# Identification of transcription factor co-binding patterns with non-negative matrix factorization

**DOI:** 10.1101/2023.04.28.538684

**Authors:** Ieva Rauluseviciute, Timothée Launay, Guido Barzaghi, Sarvesh Nikumbh, Boris Lenhard, Arnaud Regis Krebs, Jaime A. Castro-Mondragon, Anthony Mathelier

## Abstract

Transcription factor (TF) binding to DNA is critical to transcription regulation. Although the binding properties of numerous individual TFs are well-documented, a more detailed comprehension of how TFs interact cooperatively with DNA, forming either complex or co-binding to the same region, is required. Indeed, the combinatorial binding of TFs is essential to cell differentiation, development, and response to external stimuli. We present COBIND, a novel method based on non-negative matrix factorization (NMF) to identify TF co-binding patterns automatically. COBIND applies NMF to one-hot encoded regions flanking known TF binding sites (TFBSs) to pinpoint enriched DNA patterns at fixed distances. We applied COBIND to 8,293 TFBS datasets from UniBind for 404 TFs in seven species. The method uncovered already established co-binding patterns (*e.g.,* between POU5F1 and SOX2 or SOX17) and new co-binding configurations not yet reported in the literature and inferred through motif similarity and protein-protein interaction knowledge. Our extensive analyses across species revealed that 84% of the studied TFs share a co-binding motif with other TFs from the same structural family. The co-binding patterns captured by COBIND are likely functionally relevant as they harbor higher evolutionarily conservation than isolated TFBSs. Open chromatin data from matching human cell lines further supported the co-binding predictions. Finally, we used single-molecule footprinting data from mouse embryonic stem cells to confirm that the co-binding events captured by COBIND were likely occurring on the same DNA molecules.

## INTRODUCTION

The interactions between DNA and transcription factors (TFs) are crucial to transcription regulation as they intrinsically determine cell growth, development, and response to stimuli. TFs bind DNA at TF binding sites (TFBSs) in a sequence-specific manner through direct contact between DNA nucleotides and amino acids of the TFs’ DNA-binding domain (DBD) (1). Structural similarities between DBDs classify TFs into structural classes and families, and TFs from the same class or family usually recognize similar DNA patterns (or motifs).

The binding properties of individual TFs have been vastly studied (2–4), and several databases store DNA binding profiles for TFs across multiple taxonomic groups (e.g., JASPAR, CIS-BP, and HOCOMOCO (5–7)). While these databases primarily provide TF binding motifs for individual TFs, there is a need to increase our understanding of how TFs cooperatively bind DNA to regulate transcription (8). The cooperative binding of TFs generates many possible binding combinations, thus increasing the complexity of gene regulatory networks (8, 9). Recent studies argue for a flexible grammar of motifs at cis-regulatory regions, where the spacing and orientation between TFBSs would not be a key determinant for transcription regulation (10–13). Nevertheless, some TFs physically cooperate, providing a strict motif syntax important for transcription regulation. For instance, testing exhaustively the combinatorics of liver-associated TFBSs with massively parallel reporter assays revealed that TFBS orientation and order are major drivers of gene regulatory activity (14). A well-characterized example of physical interaction between TFs is POU5F1 (OCT4) cooperation with either SOX2 or SOX17 in pluripotent cells. The spacing between the binding sites will determine POU5F1’s partner and the corresponding regulative effect (15, 16). Hence, the specific spacing and orientation between the canonical motifs recognized by two TFs can characterize their co-binding at given genomic regions (17, 18). However, the cooperative binding of two TFs can also slightly modify their individual DNA sequence preference (19). Consequently, systematically identifying cooperative events genome-wide with strict binding grammar is still challenging.

The CAP-SELEX experimental technique captures co-binding events between predefined sets of TFs *in vitro* (19). However, it is still to be determined whether the same co-binding properties will necessarily occur *in vivo*. Therefore, computational methods, such as TF-COMB, TACO, SpaMo, and MCOT (20–23), leverage *in vivo* data, such as ChIP-seq (24), to predict co-binding events. These tools rely on identifying the co-occurrence of pre-defined genomic regions bound by the individual TFs or already known individual DNA binding motifs in predefined genomic regions. While these strategies reduce the search space, they restrict discoveries for the pairs of TFs where either bound regions or TF binding motifs exist for both TFs. When relying on already-known TF binding motifs, the quality of the available motif collections is a limiting factor, and this approach cannot discover new DNA sequence patterns. Another approach implemented in the RSAT *dyad-analysis* tool does not rely on pre-defined anchors or motifs and predicts spaced pairs of motifs *de novo* starting from spaced 3-mer patterns (25, 26). Finally, deep learning approaches can infer regulatory patterns from experimental data, with the capacity to pinpoint TF co-binding events (12, 27). Even though the underlying algorithms are advanced and powerful, their complexity and interpretability make it challenging to characterize specific co-binding events without extensive *a posteriori* analyses.

In this study, we aimed to discover TF co-binding patterns in the vicinity of high-quality TFBSs. The discovery of fixed co-binding patterns can be considered a matrix decomposition (or factorization) problem where an input matrix encodes nucleotides surrounding TFBSs. The ultimate goal is to group nucleotide patterns from this matrix. The non-negative matrix factorization (NMF) technique decomposes a given non-negative matrix into two low-rank, non-negative matrices to reveal underlying patterns and structures within the data (28, 29). This technique has been useful in computational biology to reveal molecular patterns from high-throughput data (28, 30). seqArchR is the first novel application of NMF for the simultaneous identification of sequence features and corresponding clusters (31). The seqArchR tool identifies critical DNA elements in promoter regions by applying NMF to the corresponding one-hot encoded sequences (31). This approach inspired us to use NMF to address the discovery of TF co-binding patterns.

We present COBIND, a Snakemake-based (32) pipeline for the *de novo* discovery of TF co-binding patterns from input sets of TFBSs (https://bitbucket.org/CBGR/cobind_tool). We applied COBIND to 8, 293 sets of high-quality TFBSs from seven species (*Arabidopsis thaliana*, *Caenorhabditis elegans*, *Danio rerio*, *Drosophila melanogaster*, *Homo sapiens*, *Mus musculus,* and *Rattus norvegicus*) for 404 unique TFs stored in the UniBind database (33). COBIND recovered established and unreported co-binding events between TFs. Among TFs from the same structural family, the majority shared co-binding patterns. In human and mouse genomes, genomic regions harboring the predicted co-binding events are evolutionarily more conserved than genomic regions without co-binding. Moreover, increased chromatin accessibility in matching human cell lines supported the predicted co-binding events from bulk experimental data. Finally, using single-molecule footprinting data from mouse embryonic stem cells, we confirmed that the predicted co-binding events between TFs likely occur on the same DNA molecules. Overall, COBIND can *de novo* discover regions in the genomes that harbor patterns of co-binding events between TFs.

## MATERIALS AND METHODS

### Predicting co-binding patterns with COBIND

COBIND takes a BED-formatted file providing the genomic coordinates of anchor regions, such as TFBS locations, as input. The tool uses NMF to reveal recurring DNA motifs with space constraints in the regions surrounding the input anchors, which are not factorized. The underlying computational pipeline consists of the following steps (Figure 1):

**Step 1: Extraction of flanking regions.** COBIND extracts the DNA sequences surrounding the anchor TFBSs (*n* bp upstream and downstream; *n=30* by default) using bedtools (v2.29.2) (34).

**Step 2: One-hot encoding.** The flanking sequences are one-hot encoded as vectors of 4 bits per DNA nucleotide (A: 1000, C: 0100, G: 0010, T: 0001; ambiguous nucleotides *∉*[ *A , C , G , T* ]) are encoded as 1111 (Figure 1 - Step 2). COBIND constructs two matrices representing the upstream and downstream sequences by combining the vectors of one-hot encoded sequences. Each row corresponds to a sequence flanking an anchor TFBS from the input set.

**Step 3: Non-negative matrix factorization.** COBIND applies NMF to each matrix using the `NMF` function of Scikit-Learn (v0.23.2) (35). The NMF decomposes an input matrix into two matrices: one representing *k* components (or factors) of nucleotide patterns and one informing the presence of the identified patterns in each row of the input matrix (28, 30). Importantly, COBIND applies NMF with multiple values of *k* to increase its capacity to capture co-binding patterns with different resolutions (see section “Parameter settings” below) (Figure 1 - Step 3). Next, COBIND assigns input sequences to each component following the methodology described by Kim and Park (36). Finally, COBIND builds motifs as positional frequency matrices (PFM) for each component by computing the occurrence frequencies of each nucleotide at each position (Additional file 1: Figure S1).

**Step 4: Motifs filtering.** COBIND filters out PFMs with information content (IC) < 2 to focus on informative patterns. Next, it aims to identify motifs corresponding to positions with local enrichment of high IC to distinguish them from motifs with scattered high IC positions or equal distribution of IC along the flanking region (Additional file 1: Figure S1). Specifically, COBIND computes the Gini coefficient *g* of each PFM to measure the inequality of IC values (37). We discard PFMs with a low Gini coefficient (*g < 0.5* by default).

**Step 5: Motif trimming.** COBIND extracts the positions corresponding to the local enrichment of high IC values revealing co-binding motifs. Specifically, it first computes the smallest window of size *n* that contains at least half of the IC of the complete flank. Finally, COBIND doubles the size of the window (adding *n/2* nucleotides up- and downstream) to define the co-binding motif.

**Step 6: Motif clustering**. COBIND can identify similar motifs multiple times since (i) NMF runs with different values for *k*, (ii) similar motifs can have different spacings with the core motif, and (iii) multiple TFBS sets can be processed in parallel, resulting in the identification of similar motifs (Figure 1 - Step 3). COBIND provides non-redundant motifs by clustering identified motifs (Figure 1- Step 6). Specifically, we use the RSAT *matrix-clustering* tool (38, 39) to cluster similar motifs (we provide the used parameters in Additional file 1: Table S1). For each motif cluster, COBIND constructs an archetypal motif following the methodology from (40), which summarizes motifs by computing the means of the counts from the aligned matrices in each cluster. We discarded from the archetypal motif the successive flanking positions with IC < 0.1.

Similarly, we cluster and summarize the anchor TFBS sequences associated with each co-binding motif (Additional file 1: Table S1).

**Step 7: Summarizing discovered co-binding events.** COBIND computes the relative spacing and orientation to the anchor motif from each co-binding archetypal motif discovered. Finally, it illustrates the corresponding co-binding events as heatmaps to visualize the co-binding motifs and the corresponding spacing and orientation to the anchor TFBSs (Figure 1 - Step 7) . The color intensity in the heatmap illustrates the proportion of input datasets with predicted co-binding motif instances. The inserted number indicates the proportion of unique sequences predicted to harbor the corresponding co-binding configurations.

**Figure 1.**
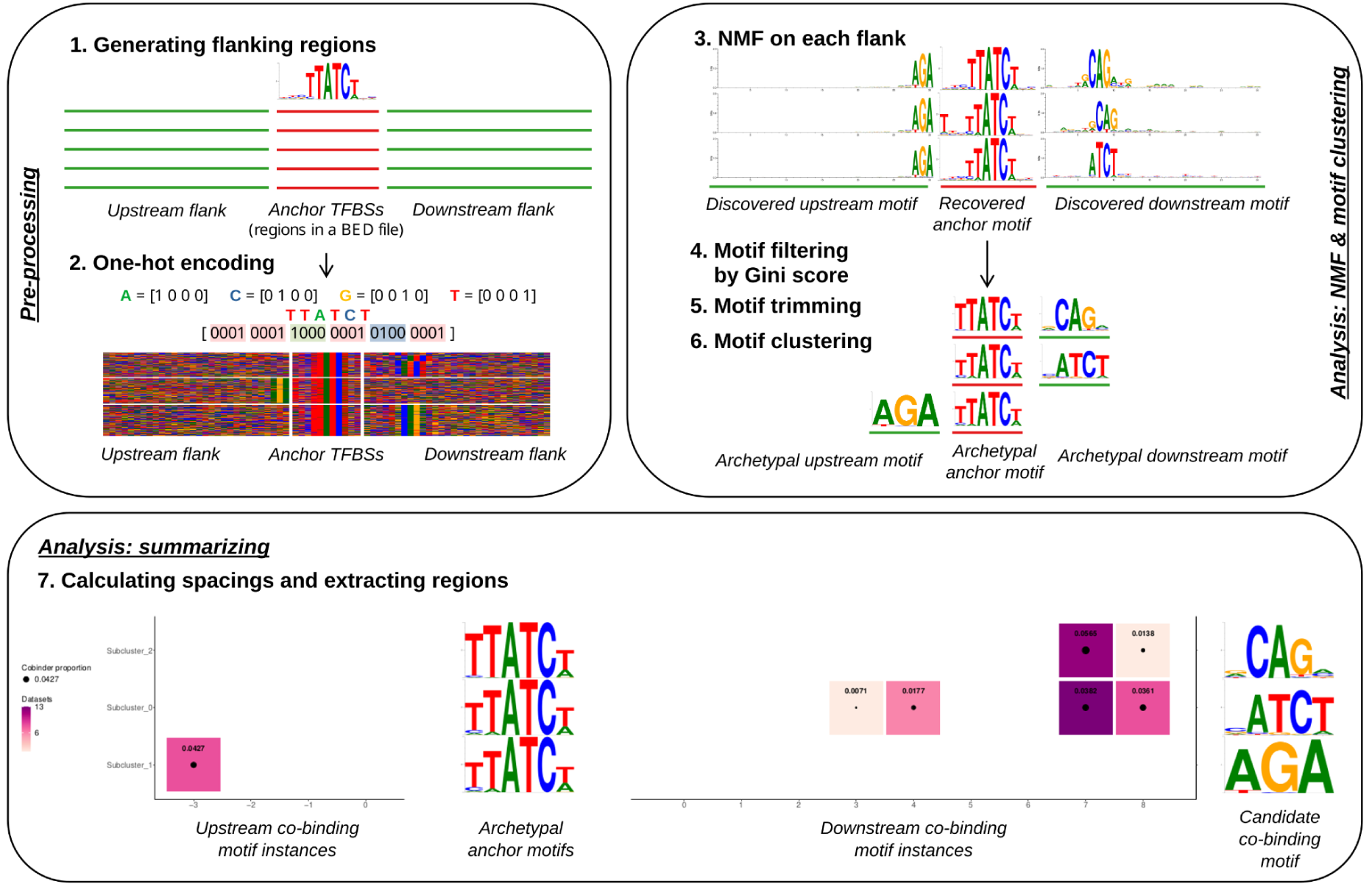
Schematic workflow of *COBIND*. **1)** COBIND takes one or multiple BED file(s) that provide the genomic location of regions of interest, such as TFBSs. The tool extracts upstream and downstream DNA sequences (default: 30 nt each). **2)** The sequences are one-hot encoded to construct encoded matrices of upstream and downstream sequences. **3)** COBIND applies NMF on each matrix to extract DNA patterns. **4)** The tool computes the Gini coefficient of each DNA pattern and discards the uninformative patterns. **5)** COBIND trims the identified flanking patterns to reveal possible TF co- binding motifs. **6)** RSAT *matrix-clustering* groups redundant motifs, and COBIND constructs archetypal motifs summarizing each cluster of similar motifs. **7)** The predicted co-binding patterns are visualized through a heatmap to illustrate each co-binding motif’s spacing, orientation, and prevalence to the anchor TFBSs.

### Parameter settings

The hyperparameters of COBIND are the length of the flanking regions, the number of components for the different runs of the NMF, and the Gini coefficient threshold. We describe below the values used in this study for each parameter and why we selected these values.

#### Length of the flanking sequences

We considered 30 bp upstream and downstream of the input TFBSs to report co-binding events likely associated with physical interactions between the TFs. The longest TFBS motif used in UniBind is about 20 bp-long (33), and cooperative TFs bind TFBSs 9- 10 bp away from each other on average (19). Furthermore, larger flanking regions resulted in noisier co-binding motifs and could miss co-binding motif configurations (Additional file 1: Figure S2).

#### Number of components

COBIND runs NMF several times with different numbers of components (*k*). We used synthetic data to estimate a suitable range of values for *k*. Specifically, we implanted different proportions of instances of predefined motifs in random sequences. We considered sets of 1000, 5000, 10, 000, 50, 000, and 100, 000 random sequences. For each set of random sequences, we injected an instance of the considered motif in 0.5, 1, 2, 3, 4, 5, and 6% of the sequences. We ran COBIND with k ∊ [2, 17] for each synthetic dataset. We assessed specificity and sensitivity by counting the number of correct motifs predicted by COBIND for each value of *k*.

Moreover, we reported the proportion of sequences correctly predicted to contain the injected motifs. We considered that COBIND predicted the correct motif if it was similar to the injected motif - as assessed by Tomtom (41) with a p-value < 0.05 (Additional file 1: Table S1). We used the results from the synthetic data to select *k* ∊ [3, 6] for further analyses (Additional file 1: Figure S3 for an example).

#### Gini coefficient threshold

To empirically determine the Gini coefficient threshold, we compared the distributions of Gini coefficients for the motifs predicted by COBIND on the TFBS datasets from UniBind (see below) with those of predictions from random sequences. For each UniBind set of TFBSs, we constructed a set of random sequences by shuffling the DNA sequences flanking the TFBSs with the *kmer shuffling* module of the *BiasAway* tool with *k=1* (42). For each species considered, we selected the Gini coefficient threshold corresponding to a 1% false discovery rate (Additional file 1: Figure S4). This strategy resulted in the following Gini coefficient thresholds: 0.51 for *Arabidopsis thaliana*, 0.45 for *Caenorhabditis elegans*, 0.52 for *Danio rerio*, 0.48 for *Drosophila melanogaster*, 0.51 for *Homo sapiens*, 0.53 for *Mus musculus*, and 0.52 for *Rattus norvegicus*. Finally, we estimated the stability of the thresholds obtained with this strategy by performing the same analysis multiple times and subsampling the number of datasets from *H. sapiens* and *M. musculus*. Overall, we observed that the Gini coefficient thresholds were stable across the different runs (Additional file 1: Figure S5).

### Transcription factor binding site datasets

We used the robust collection of TFBS sets from the UniBind database (2021 version) (https://unibind.uio.no/) derived from 7 species (*Arabidopsis thaliana*, *Caenorhabditis elegans*, *Danio rerio*, *Drosophila melanogaster*, *Homo sapiens*, *Mus musculus, Rattus norvegicus*) (33). UniBind TFBS datasets derive from pairs of ChIP-seq peak datasets and JASPAR TF binding profiles; some TFs are associated with several binding profiles. Datasets with less than 1000 TFBSs were filtered out (Additional file 1: Table S2).

### Benchmarking

#### Synthetic datasets

We generated the synthetic datasets following the methodology mentioned above. We constructed synthetic datasets with 500, 1000, 3000, 5000, 10, 000, and 50, 000 sequences with instances of a known motif inserted in 0.5, 1, 3, 5, and 10% of the sequences. We considered the motif associated with the ATF4 TF (MA0833.2) in the JASPAR database (7). To run SpaMO, we inserted the CTCF (MA0139.1) motif at the center of the flanking sequences since SpaMO requires a motif to anchor its search for co-binding patterns.

#### Comparisons between COBIND and other tools

We used the synthetic data to compare COBIND with MEME-ChIP (43), RSAT peak-motifs *dyad-analysis* (25, 26, 44, 45), RSAT peak-motifs *oligo-analysis* (26, 44, 45), RSAT peak-motifs *position-analysis* (44, 45), and SpaMo (21). We compared the motifs predicted by the different tools with the original inserted motifs using Tomtom (41) (Additional file 1: Table S1). We considered a discovered motif as a match to the injected motif if Tomtom predicted them to be similar with a p-value < 0.05. To run SpaMo, we used the JASPAR 2020 CORE collection of non-redundant profiles (46). In addition, we computed the observed over the expected number of sequences predicted to contain instances of the predicted motif. We expected the optimal observed/expected ratio to be within [0.5- 1.5] to balance specificity and sensitivity. A smaller ratio indicates that the discovery algorithm missed more than half of the injected sites. A larger ratio indicates that the discovery algorithm predicted 1.5 times more injected sites than expected.

We ran all tools with their default parameters. When estimating the running time, we used the parallelizable version of the NMF for COBIND.

### Inference of the co-binding transcription factors from the discovered co-binding motifs

We assessed the similarities between a predicted co-binding motif and known motifs associated with 5, 272 TFs - collected from JASPAR 2022 non-redundant CORE, JASPAR 2022 non-redundant Unvalidated taxon-specific, and CIS-BP (5, 7) - using Tomtom (Additional file 1: Table S1) (41). We predicted TF_B_ to bind the predicted co-binding motif associated with anchor TFBSs of TF_A_ if Tomtom reported a similarity p-value < 0.05 with the known canonical motif bound by TF_B_ (criteria I). To assess possible physical interactions between TF_A_ and TF_B_, we retrieved protein-protein interaction (PPI) data from STRING v11.0 (47). We considered TF_A_ and TF_B_ to interact physically when their STRING PPI score was above or equal to 500 (criteria II). We reported the co-binding pair TF_A_-TF_B_ to be already “known” when they met criteria I and II. We reported TF_A_-TF_B_ as a novel co-binding pair when they met criteria I but not II.

### Shared co-binding motifs across transcription factor structural families

To assess the conservation of co-binding motifs across TFs, we ran COBIND (up to Step 5) on all UniBind datasets, excluding UniBind predictions derived from dimer anchor JASPAR TF binding profiles. We independently clustered all the predictions (COBIND Step 6) for each TF structural family; note that this step considered TFs from multiple species. When the co-binding motifs associated with two (or more) TFs from the same family were similar (i.e., belonged to the same cluster), we considered the TFs to share the same co-binding motif.

### Evolutionary conservation

We downloaded the following conservation tracks in bigwig format from the UCSC genome browser: phastCons100way (human; hg38), phastCons60way (mouse; mm10), phastCons135way (*C. elegans*; WBcel235), and phastCons27way (*D. melanogaster*; dm6) (48). We obtained the 63 flowering plants PhastCons conservation track from PlantRegMap (49) for analyzing datasets from *A. thaliana*. We computed the conservation scores of a genomic region using the *aggregate* and *extract* functions of bwtool (v1.0) (50). We compared the distributions of conservation scores between two regions with a one-sided Wilcoxon signed-rank test.

### DNase I hypersensitive footprinting analysis

We downloaded DNase I hypersensitivity (DHS) data generated by the ENCODE consortium for multiple human cell types in bigwig format at https://resources.altius.org/~jvierstra/projects/footprinting.2020/ (51). We considered the 53 cell lines or tissues for which both DHS and COBIND predictions were available; we analyzed 66 DHS datasets associated with 87 co-binding configurations for 56 TF binding profiles as anchors. We followed the methodology described above to compare evolutionary conservation scores when comparing the DHS signal between two genomic regions (with or without a co-binding event predicted by COBIND).

### Single-molecule footprinting analysis

We downloaded 12 single-molecule footprinting (SMF) datasets derived from triple- methyltransferases knockout experiments in mouse embryonic stem cells (mESCs) from ArrayExpress with accession numbers E-MTAB-9033 and E-MTAB-9123 (52). Whenever applicable, we performed the analyses of SMF data with the SingleMoleculeFootprinting R package following the instructions previously described (53). To compare the SMF data in the genomic regions predicted by COBIND to contain or not a co-binding event in mESC datasets, we modified the SortReadsByTFCluster_MultiSiteWrapper functions from the development version of the SingleMoleculeFootprinting package (54) (https://bitbucket.org/CBGR/cobind_manuscript/src/master/bin/single_molecule/analyse_sm/sort_tf_reads_by_cobind_clusters.R). Using the methylation status provided by the SMF data, we determined each genomic region to be either (i) free of nucleosomes (“Accessible”), (ii) occupied by nucleosomes (“Nucleosome”), (iii) bound only by the anchor (“Anchor”), (iv) bound only by the co-binding TF (“Co-binding”), or (v) bound by both the anchor and co-binding TF (“Anchor + Co-binding”). We summed the number of molecules for each state in different samples and computed the proportions of molecules. We compared the proportions of each state in genomic regions predicted to contain a co-binding event or not using a two-sided Wilcoxon signed-rank test.

## RESULTS

### COBIND discovers co-binding patterns *de novo*

#### Comparisons with other tools on simulated data

We report a new computational framework, COBIND, to predict co-binding patterns in the vicinity of TFBSs provided as input. COBIND relies on applying NMF to the one-hot encoded regions flanking the TFBSs to predict co-occurring DNA motifs with fixed spacing (Materials and Methods). We compared COBIND to other tools discovering motifs *de novo* (MEME-ChIP, RSAT *dyad-analysis*, RSAT *oligo-analysis*, and RSAT *position-analysis*). Additionally, we compared COBIND to SpaMO, which predicts spatially co-occurring instances of known motifs (Additional file 1: Table S3). For comparisons, we generated synthetic data by injecting instances of a known motif at different frequencies in random sets of 500 to 50, 000 DNA sequences (Materials and Methods).

COBIND predicted the correct motif inserted when considering a minimum of 1000 sequences, with at least 3% of the sequences containing instances of the injected motif (Additional file 1: Figure S6). Compared to the other tools, COBIND predicted no false positive motif across most configurations (see the number of motifs discovered that were not similar to the injected motif in Additional file 1: Figure S6). While SpaMo discovered the correct inserted motif in most cases, it came at the cost of predicting many false positives. One should note that SpaMo requires a reference set of known motifs to predict co-occurrences, while COBIND predicts co-binding motifs *de novo*. Furthermore, we found that COBIND predicted a more significant proportion of sequences harboring the injected motif more similar to the expected number than the other tools (see the observed over expected ratios in Additional file 1: Figure S6). COBIND was slower than other tools with large data sets (>50, 000 sequences) due to the motif clustering step applied to the results of the NMF with 3 to 6 components. Nevertheless, COBIND was faster on datasets with < 50, 000 sequences (∼500 seconds on 50, 000 sequences) as it allows for parallelization across CPU cores (Additional file 1: Figure S7).

#### Proof-of-concept: POU5F1 co-binding with SOX2 or SOX17

As a proof-of-concept, we applied COBIND to UniBind TFBS datasets for the SOX2 and SOX17 TFs. Previous studies established that both TFs partner with POU5F1 in pluripotent cells, where POU5F1 co-binds with SOX2 at regulatory elements to promote pluripotency. At the same time, it co-binds with SOX17 to control endoderm differentiation at other regulatory elements (11, 16). While POU5F1 and SOX2 cooperate through binding at instances of their respective canonical motifs, POU5F1 can associate with SOX17 to bind a compressed motif (Figure 2A). COBIND successfully discovered the two co-binding patterns - canonical and compressed - recognized by POU5F1 when applied to the SOX2 and SOX17 TFBS datasets from *H. sapiens* and *M. musculus* (Figure 2B). COBIND retrieved the correct pattern in 12 human SOX2 datasets (58%), with ∼7% of the sequences predicted to harbor the pattern, and in 24 mouse datasets (32%), with ∼6% of the sequences harboring the pattern (Additional file 1: Figure S8A-B). In agreement with previous studies, the datasets where COBIND predicted the pattern derived from human and mouse ESC and mouse embryonic fibroblast (16, 18, 55). When considering the SOX17 datasets, COBIND successfully discovered the canonical co-binding pattern and the compressed motif. It predicted the compressed co-binding pattern in 2 datasets (50%; derived from mouse ESC), with 5% of the sequences harboring the co-binding motif (Additional file 1: Figure S8C).

**Figure 2.**
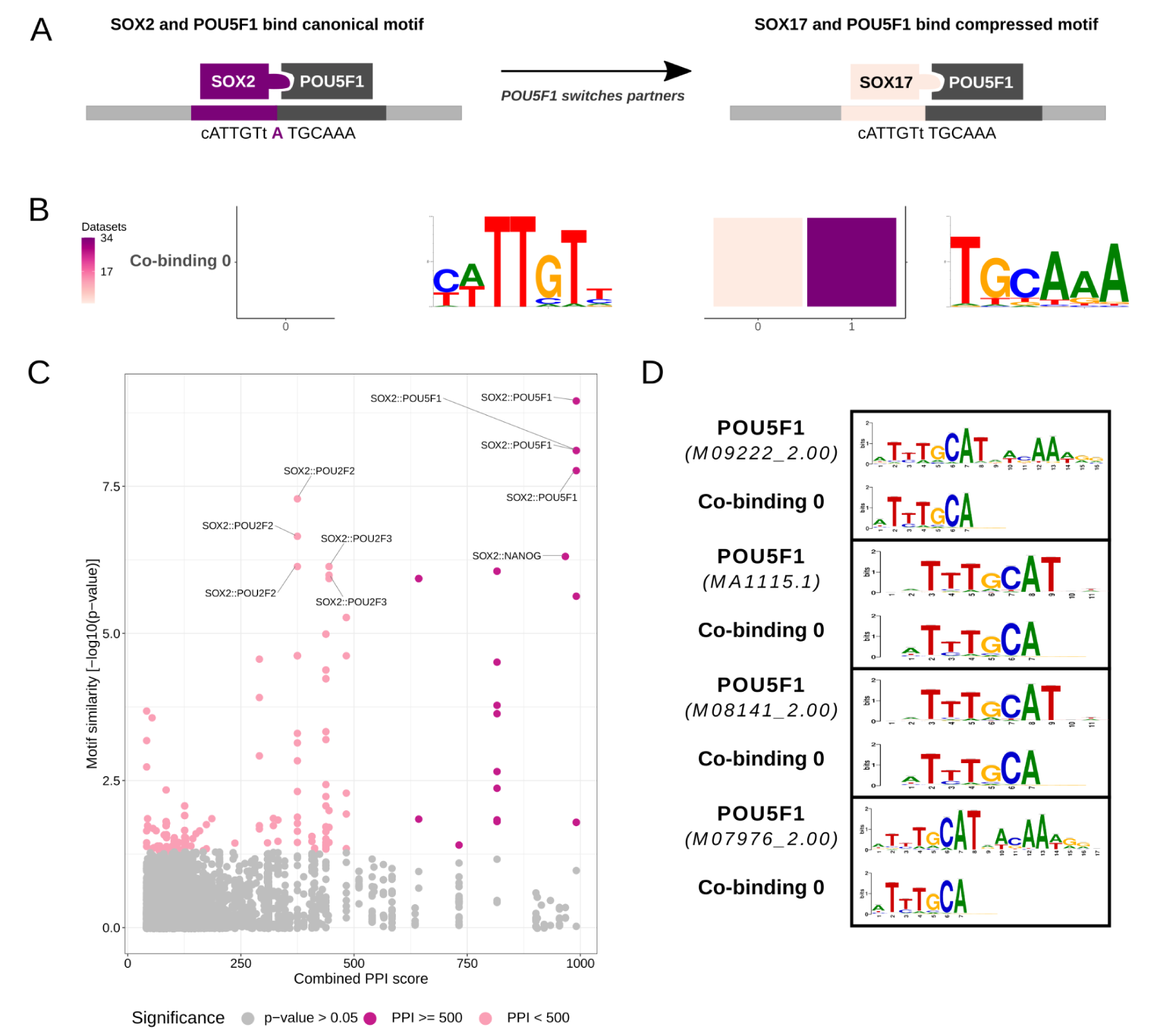
Application of COBIND to SOX2 and SOX17 TFBS datasets. COBIND predicts POU5F1 to co-bind with SOX2 (human and mouse) and SOX17 (mouse), upholding the previously known mechanism of POU5F1 switching partner TFs and binding motifs with different syntax. **A**. POU5F1 partners with SOX2 to bind its canonical motif. In contrast, POU5F1 associates with SOX17 to bind a compressed motif. **B.** The co-binding patterns discovered in SOX2 and SOX17 datasets correspond to the expected canonical and compressed motifs. **C**. The co-binding motif found in the SOX2 datasets was associated with POU5F1 using motif similarity and PPI data (significant motif similarity p-value < 0.05 and PPI combined score > 500). **D.** Visual representation of the similarity between the discovered co-binding motif and motifs recognized by POU5F1 in JASPAR and CIS-BP.

In this proof-of-concept example, we knew *a priori* that POU5F1 was the binding partner of SOX2 and SOX17. Nevertheless, COBIND discovered *de novo* the co-binding motif. In other settings, one would not know the TF binding to the co-binding motif discovered. To address this challenge, we combined motif similarity to already known motifs from JASPAR and CIS-BP with protein-protein interaction (PPI) data from the STRING database to infer the TFs potentially binding the motifs revealed by COBIND (Materials and Methods). This strategy confirmed that the framework inferred POU5F1 as the binding partner of SOX2 and SOX17 (Figure 2C-D; Additional file 1: Figure S8).

#### COBIND discovers known and novel co-binding patterns in a large-scale analysis across several species

We ran COBIND on 8, 293 UniBind TFBS datasets associated with 404 unique TFs from 7 species (Materials and Methods; Additional file 1: Table S2). Beyond predicting co-binding patterns with COBIND, we inferred the TFs likely binding to the discovered patterns following the strategy described above (also see Materials and Methods). Altogether, COBIND revealed 898 co-binding patterns for 218 TFs (22 in *A. thaliana* datasets, 7 in *C. elegans*, 4 in *D. rerio*, 22 in *D. melanogaster*, 420 in *H. sapiens*, 407 in *M. musculus*, and 16 in *R. norvegicus*). For half of the reported co-binding motifs (450 out of 898), we found motif similarity and PPI data to support the inferred pair of co-binding TFs (Figure 3). Specifically, the data supported all co-binding patterns for *D. rerio* and *C. elegans*, more than half of the patterns for *R. norvegicus*, *H. sapiens*, and *M. musculus*, 18.2% for *A. thaliana*, and 31.8% for *D. melanogaster*. Furthermore, the analysis of motif similarity between COBIND’s predictions revealed that 84% (163 out of 193) of the TFs shared a co-binding motif with another member of the same TF structural family (Materials and Methods), which confirmed the conservation of co-binding patterns within families of TFs across species. We provide all predictions for the community to explore through a dedicated website at https://cbgr.bitbucket.io/COBIND_2023/.

**Figure 3.**
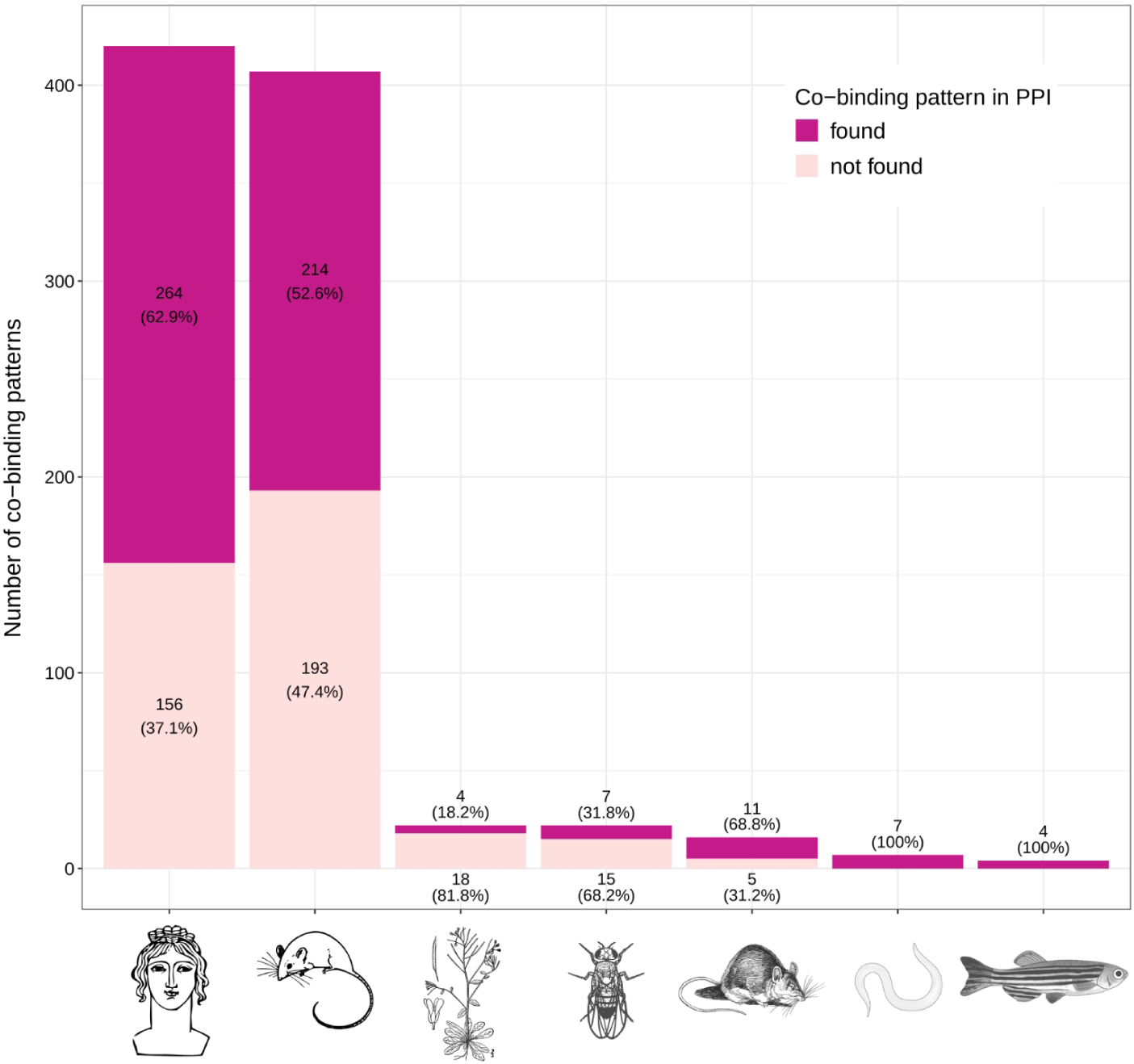
Overview of COBIND predicted co-binding patterns. The bar plot provides the number of co-binding patterns (y-axis) discovered by COBIND with (dark purple) or without (light purple) support from motif similarity and PPI data. The bars contain the corresponding numbers and percentages. Each bar summarizes the results for a species (x-axis).

As a case example, COBIND predicted two co-binding patterns around TEAD1 TFBSs. Specifically, COBIND revealed the same motif with different spacings from the provided anchor TFBSs (2nt upstream or 3nt downstream of TEAD1 TFBSs; (Additional file 1: Figure S9A)). We found that the two co-binding patterns occurred in ∼7% of the input sequences (2 bp upstream pattern) from mouse nerve tumor cells and in ∼3% of the sequences from human pancreas and astrocytoma cells (3 bp downstream pattern). Our approach inferred that TEAD2 and TEAD4 cooperate with TEAD1 (Additional file 1: Figure S9B-C). Previous studies reported that TEAD TFs bind either an isolated M-CAT element or direct DNA repeats with spacing ranging from 0 to 6 bp (56–58). These studies further support the co-binding patterns predicted by COBIND. The predictions around TEAD1 TFBSs provide an example of COBIND’s capacity to predict co-binding events supported by both motif similarity and PPI data.

To exemplify predictions of new co-binding TFs, COBIND revealed a co-binding motif located 4 bp upstream of HY5 anchor TFBSs in *A. thaliana* (Additional file 1: Figure S10A). The analyses of TFBS datasets associated with the TFs ABF1, ABF3, ABF4, GBF2, and GBF3 displayed the same co-binding pattern. We found that the canonical motifs recognized by human NF-Y TFs (NF-YC and NF-YA) were similar to the co-binding motif discovered by COBIND (Additional file 1: Figure S10B-C). However, no PPI data currently support the physical interactions between the anchor TFs and NF-Y orthologs in plants. Nevertheless, the bZIP67 TF, which belongs to the same basic leucine zipper (bZIP) family as HY5, ABF1, ABF3, ABF4, GBF2, and GBF3, is known to form a transcriptional complex together with NF-YC2 and bind ER stress response elements (ERSEs) to regulate omega-3 fatty acid content in *A. thaliana* (59). In agreement with this observation, we noted that the anchor motif combined with the co-binding motif discovered by COBIND forms the motif associated with ERSEs (60). Furthermore, HY5 competes with bZIP28, another member of the bZIP family, to bind ERSEs on promoters of unfolded protein response genes, and bZIP28 interacts with NF-Y protein complexes (60, 61). Consistent with this knowledge, we found that most of the genomic regions predicted by COBIND to harbor the co-binding events in the HY5 dataset were in promoter regions (Additional file 1: Figure S10D). Altogether, these multiple lines of evidence strongly support the co-binding patterns predicted by COBIND between bZIP and NF-Y TFs in *A. Thaliana*.

### COBIND discovers the extended motif bound by CTCF

We investigated another example where COBIND predicted a co-binding pattern unsupported by motif similarity and PPI data. Specifically, COBIND identified a co-binding motif in the proximity of CTCF TFBSs for several datasets (Figure 4A). When analyzing human and mouse datasets, we observed two spacing configurations between the anchor CTCF TFBSs and the predicted co-binding events (Additional file 1: Figure S11A-B). When considering datasets from zebrafish, COBIND revealed only one spacing configuration (Additional file 1: Figure S11C). Across the datasets, COBIND predicted from 1.4 to 4% of the sequences to harbor the co-binding patterns. All datasets from the human and 23% of the mouse datasets (87 out of 382) contained the identified co-binding patterns. Previous studies have reported the motif predicted by COBIND as an extension of the canonical CTCF motif (62, 63). These studies revealed that the eighth zinc finger of CTCF acts as a linker instead of a clamp to allow zinc fingers 9-11 to bind the extended motif (Figure 4B), affecting the binding efficiency, residence time, and binding off-rate of CTCF (62, 63). Consequently, COBIND did not predict co-binding between distinct TFs but an extended motif seldomly bound by CTCF through different combinations of zinc finger contacts with the DNA.

**Figure 4.**
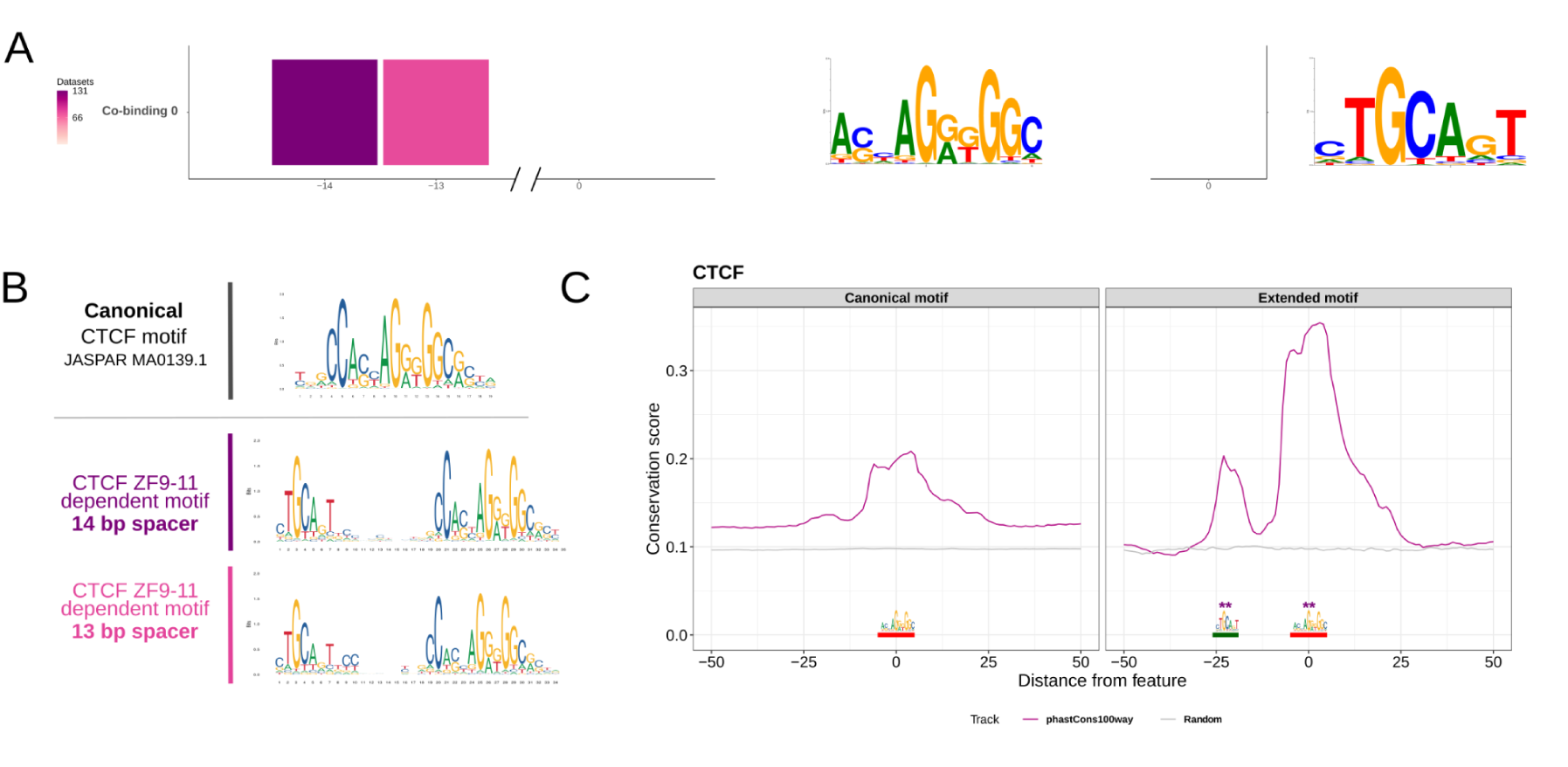
CTCF binds a conserved extended motif. (**A**) COBIND predicted two binding configurations through an extended motif for CTCF in human and mouse datasets. (**B**) The two binding configurations derive from contacts between the zinc fingers (ZF) 9-11 and the DNA at the upstream motif. (**C**) Comparison between vertebrate evolutionary conservation of genomic regions harboring the CTCF canonical motif (left) and regions with the extended motif revealed higher conservation of the extended motif. The purple lines provide the mean conservation score across the regions considered. The gray lines provide the mean conservation score across the same number of random genomic regions in the human genome. ** indicates a Wilcoxon test p-value < 0.001. The anchor CTCF motif is underlined in red, and the upstream portion of the extended motif is underlined in green.

Since evolutionary conservation is a hallmark of functional importance (48), we assessed the functional relevance of the genomic regions harboring the discovered binding patterns by examining their evolutionary conservation across vertebrates (Materials and Methods). We observed that the regions containing the extended CTCF motif were more conserved than regions with the canonical motif (Wilcoxon test p-value < 0.001; Figure 4C). The increased evolutionary conservation was consistent across all spacing configurations observed in human, mouse, and zebrafish. These results exemplify how COBIND can predict relevant DNA patterns that do not correspond to the co-binding of two proteins but to binding variants for the same TF, a particular case of the zinc finger families.

### Genomic regions harboring a co-binding pattern are evolutionarily more conserved than regions without co-binding

We assessed the functional relevance of all the co-binding patterns discovered by COBIND across species by analyzing their evolutionary conservation (Materials and Methods). Specifically, we compared the evolutionary conservation of the genomic regions harboring a co-binding pattern, named co-bound regions for simplicity, with those without a predicted co-binding pattern. We first describe the analyses of the case studies presented above as examples. We found that the co-bound regions associated with SOX2::POU5F1 and TEAD1::TEAD in the human and mouse genomes exhibited a significantly increased evolutionary conservation compared to those without predicted co-binding (Wilcoxon test p-value < 0.001; Figure 5A; Additional file 1: Figure S12A). Notably, we observed increased conservation at the co-binding and the anchor motifs. When considering the co-binding pattern associated with HY5 in *A. thaliana*, we did not observe a significant difference in conservation between the regions with and without the co-binding pattern. Nevertheless, both the anchor and the co-binding motifs exhibited peaks of evolutionary conservation, confirming the likely functional relevance of the predicted co-binding pattern across the flowering plant kingdom (Additional file 1: Figure S13A).

**Figure 5.**
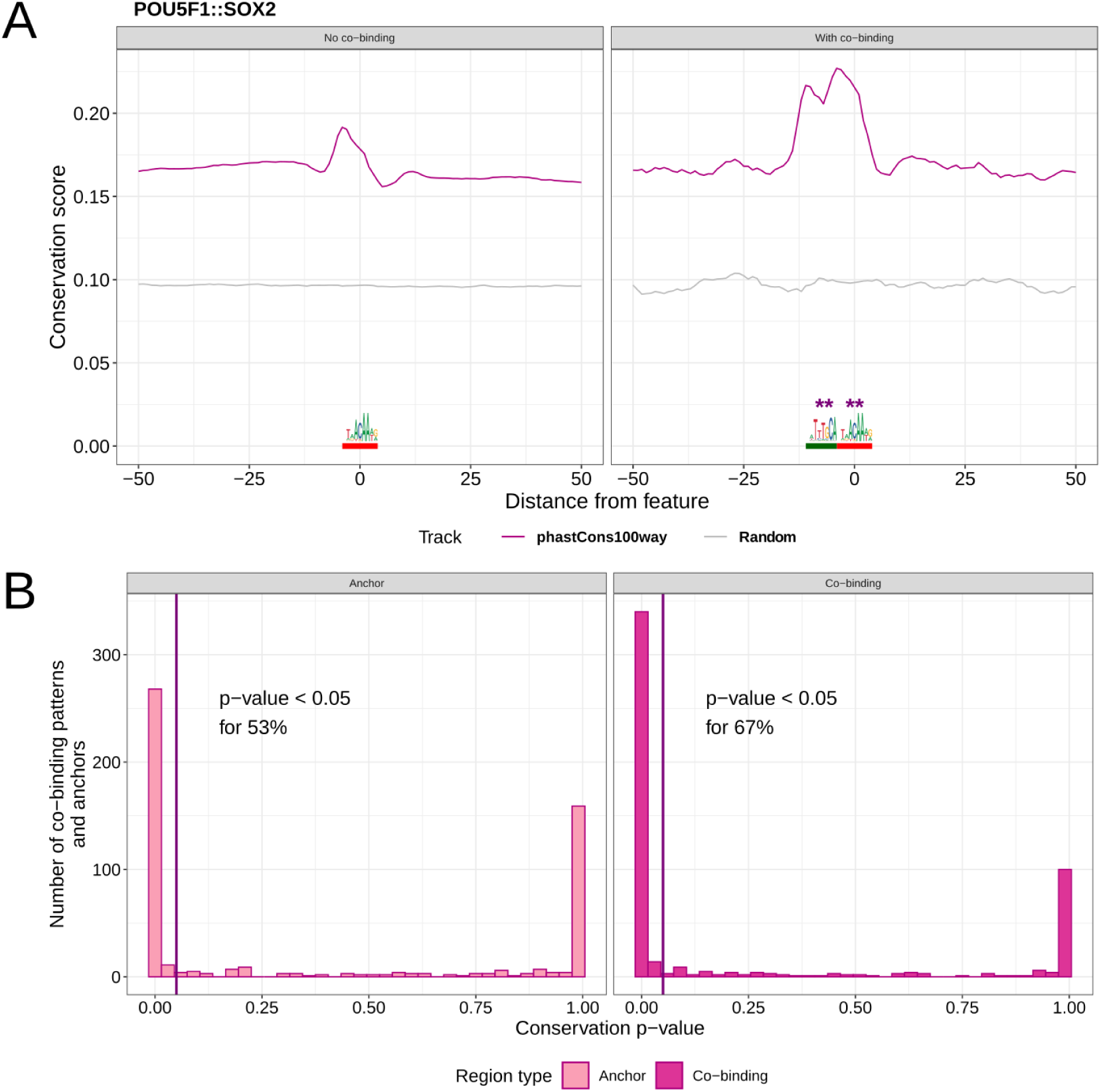
Analysis of the evolutionary conservation of co-binding patterns discovered by COBIND in the human genome. (**A**) Comparison between vertebrate evolutionary conservation of genomic regions where COBIND predicted a co- binding pattern (right) or not (left). The purple lines provide the mean conservation score across the regions considered. The gray lines provide the mean conservation score across the same number of random genomic regions in the human genome. ** indicates a Wilcoxon test p-value < 0.001. The anchor SOX2 motif is underlined in red, and the co-binding motif is underlined in green. (**B**) We compared the evolutionary conservation scores at the anchors’ motifs in genomic regions harboring a co-binding pattern or not. The left panel represents the histogram of the number of co-binding patterns where the evolutionary conservation was higher in genomic regions harboring a co-binding pattern. The right panel represents the corresponding histogram when comparing genomic locations of the co-binding motif.

We systematically compared the evolutionary conservation at instances of the anchor and co-binding motifs discovered by COBIND across all datasets and species. Altogether, we observed across species that 14% (for *A. thaliana*), 42% (for *D. melanogaster*), 50% (for *C. elegans*), 67% (for *H. Sapiens*), and 69% (for *M. musculus*) of the co-binding patterns exhibited significantly more conservation of the co-binding motifs in co-bound regions than in the other regions (Figure 5B; Additional file 1: Figure S12B-S14). When considering the anchor motifs solely, we found that they were more conserved in the co-bound regions for 32% (*A. thaliana*), 38% (for *C. elegans*), 50% (for *D. melanogaster*), 52% (for *M. musculus*), and 53% (*H. sapiens*) of the datasets (Figure 5B; Additional file 1: Figure S12B-S14). The consistent increased evolutionary conservation of the co-binding patterns supports the functional importance of COBIND’s predictions across species. Furthermore, our results suggest that the underlying genomic regions harboring a fixed binding motif syntax are evolutionarily important.

### DNase I hypersensitive footprints support the predicted co-binding patterns

We further assessed the co-binding patterns discovered by COBIND by analyzing orthogonal experimental data probing chromatin openness. The DNase-seq assay captures open chromatin regions by revealing DNase I hypersensitive sites (DHS) (64). Importantly, TFs interacting with the DNA in open chromatin regions protect their TFBSs from the DNase cleavage, which leaves a footprint on the corresponding TFBSs (51, 65). We retrieved 66 DHS footprint datasets from 53 human cell types (51) that matched some of the UniBind TFBS datasets used in this study. This data allowed us to investigate DHS footprints at the discovered co-binding motifs for 87 co-binding patterns associated with 56 TF binding profiles as anchors. For each co-binding pattern, we compared the depth of the DHS footprint at the locations of the co-binding motifs between the co-bound regions and the other regions (Materials and Methods).

As we ran COBIND on regions surrounding TFBSs predicted as high-quality direct TF-DNA interactions with ChIP-seq and computational evidence from UniBind, we expected DHS footprints at the anchor TFBSs. Indeed, we observed footprints of TF-DNA interactions; for instance, the DHS footprints observed for the CTCF datasets (Figure 6A). Furthermore, the analyses exhibited deep DHS footprints at the location of the co-binding motifs. Overall, we found significantly deeper DHS footprints at the locations of the co-binding motifs in the co-bound regions than in the other regions for 83% of the pairs of co-binding patterns - DHS dataset (Figure 6B). Notably, the anchor TFBSs at co-bound regions exhibited deeper DHS footprints than the TFBSs in other regions for 81% of the pairs (Figure 6B). Co-bound and anchor TFBS locations showed deeper DHS footprints in the genomic regions predicted with a co-binding pattern for 76% of the pairs (associated with 27 TFs - 61%). As CTCF datasets represented 43% of the total datasets analyzed here, we performed the same analysis excluding the CTCF datasets. We found that 45% of the co-binding pattern - DHS dataset pairs showed significant DHS footprints at the anchor TFBSs, and 57% exhibited significantly deeper DHS footprints at the co-binding motif locations in co-bound regions than in the other regions. Altogether, 38% of the pairs exhibited deeper DHS footprints at both the co-bound and anchor TFBS locations. Overall, the DHS footprint analyses supported the co-occupancy of the anchor and the co-binding motifs at the co-bound regions.

**Figure 6.**
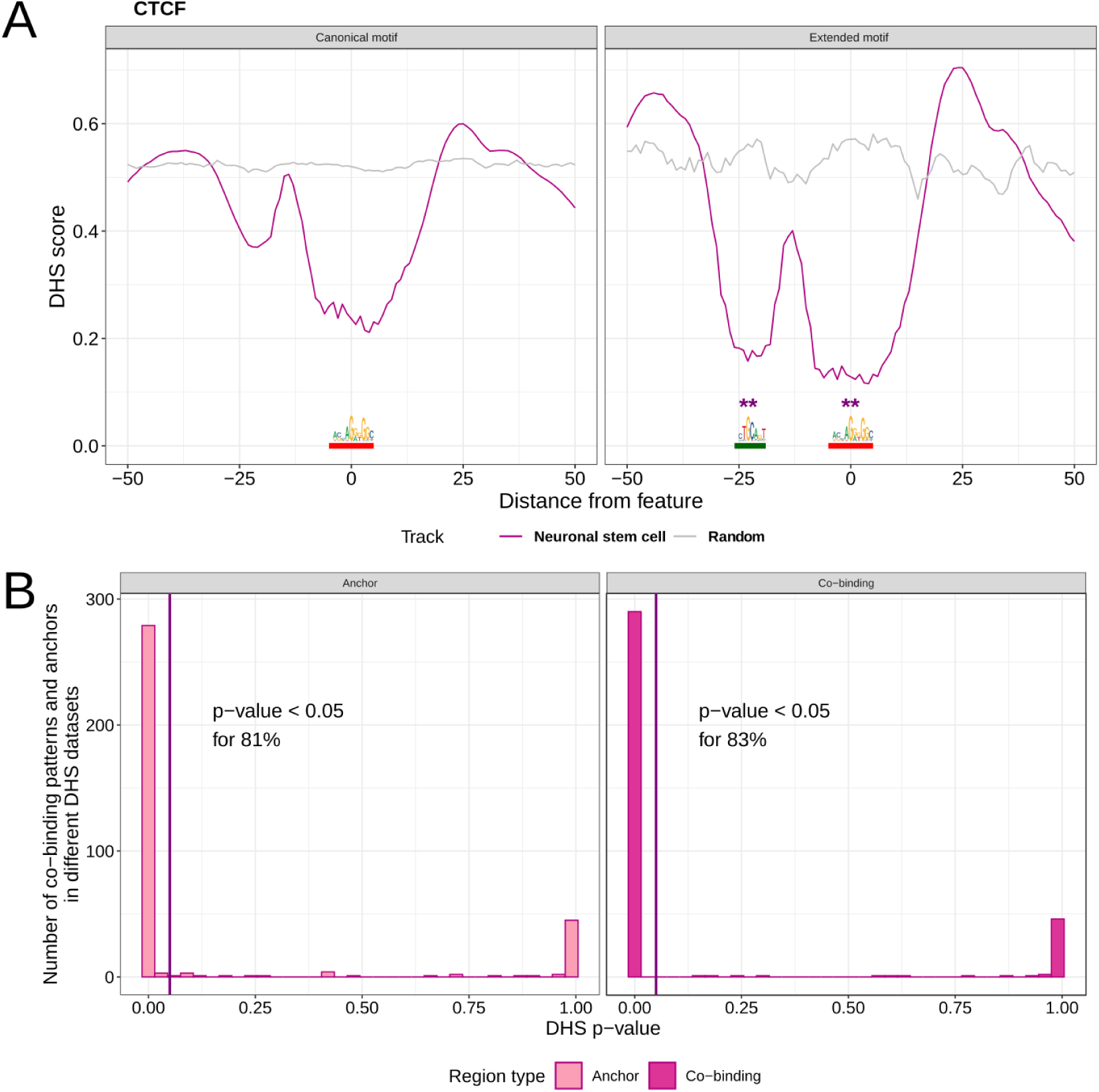
DHS footprinting analyses at anchor and co-binding locations. (**A**) The plots represent the average DHS score in regions surrounding CTCF TFBSs in neuronal stem cells (purple lines) or at random regions (gray lines). The left plot provides DHS scores at genomic regions where COBIND did not predict the CTCF extended motif; the right plot provides the DHS scores at genomic regions harboring the CTCF extended motif. ** indicates a Wilcoxon test p-value < 0.001. The anchor CTCF motif is underlined in red, and the upstream portion of the extended motif is in green. (**B**) We compared DHS scores at the anchors’ motifs in genomic regions harboring a co-binding pattern or not. The left panel represents the histogram of the datasets where the DHS footprint was deeper in genomic regions harboring a co-binding pattern. The right panel represents the corresponding histogram when comparing genomic locations of the co-binding motif.

### Single-molecule footprints support TF co-occupancy at single-molecule resolution

The DHS footprinting analysis described above relied on bulk DNase-seq data. Consequently, the results provided average estimations of the co-occupancy of TFs across cells. We aimed to investigate the co-occupancy of TFs at the co-binding patterns predicted by COBIND at single-molecule resolution. To this end, we considered single-molecule footprinting (SMF) data, which probed the co-occupancy of TFs and nucleosomes at single-molecule resolution for accessible genomic regions in mouse embryonic stem cells (mESC) (52). The SMF data overlapped 3, 373 co-bound regions (909, 918 reads) and 94, 387 regions (24, 015, 952 reads) not predicted to contain a co-binding pattern for 16 TFs in mESC (Materials and Methods).

Figure 7A-B illustrates the SMF data analysis at two genomic regions predicted to harbor co-binding events for SOX2 and POU5F1 (Figure 7A) and the extended CTCF motif (Figure 7B). For each region, we determined the fractions of molecules predicted as (i) accessible, (ii) occupied by nucleosomes, (iii) only occupied at the anchor motif, (iv) only occupied at the co-binding motif, or (v) co-occupied at both the anchor and the co-binding motifs. In the two example regions, we observed footprints of co-occupancy at both the anchor motif and the co-binding motif locations. Specifically, the analysis of 273 molecules across five replicates revealed co-occupancy for 27% of the molecules when considering the SOX2-associated region (Figure 7A). Five hundred ninety-two molecules across six replicates revealed co-occupancy of the extended CTCF motif for 59% of the molecules (Figure 7B). When considering all the regions with SMF data, we found a larger fraction of co-occupied anchor and co-binding motif locations on the same molecule in COBIND-predicted co-bound regions than in other regions (p-value < 0.001; Figure 7C). However, not all anchor TF binding profiles satisfied this observation (Additional file 1: Figure S15). Moreover, we found that genomic regions predicted by COBIND to harbor a co-binding pattern were significantly more occupied (for all occupancy states independently) than the other genomic regions (p-value < 0.001; Figure 7C). Altogether, the results confirmed that the genomic regions predicted by COBIND to harbor a co-binding pattern were overall more accessible or co-occupied at single-molecule resolution.

**Figure 7.**
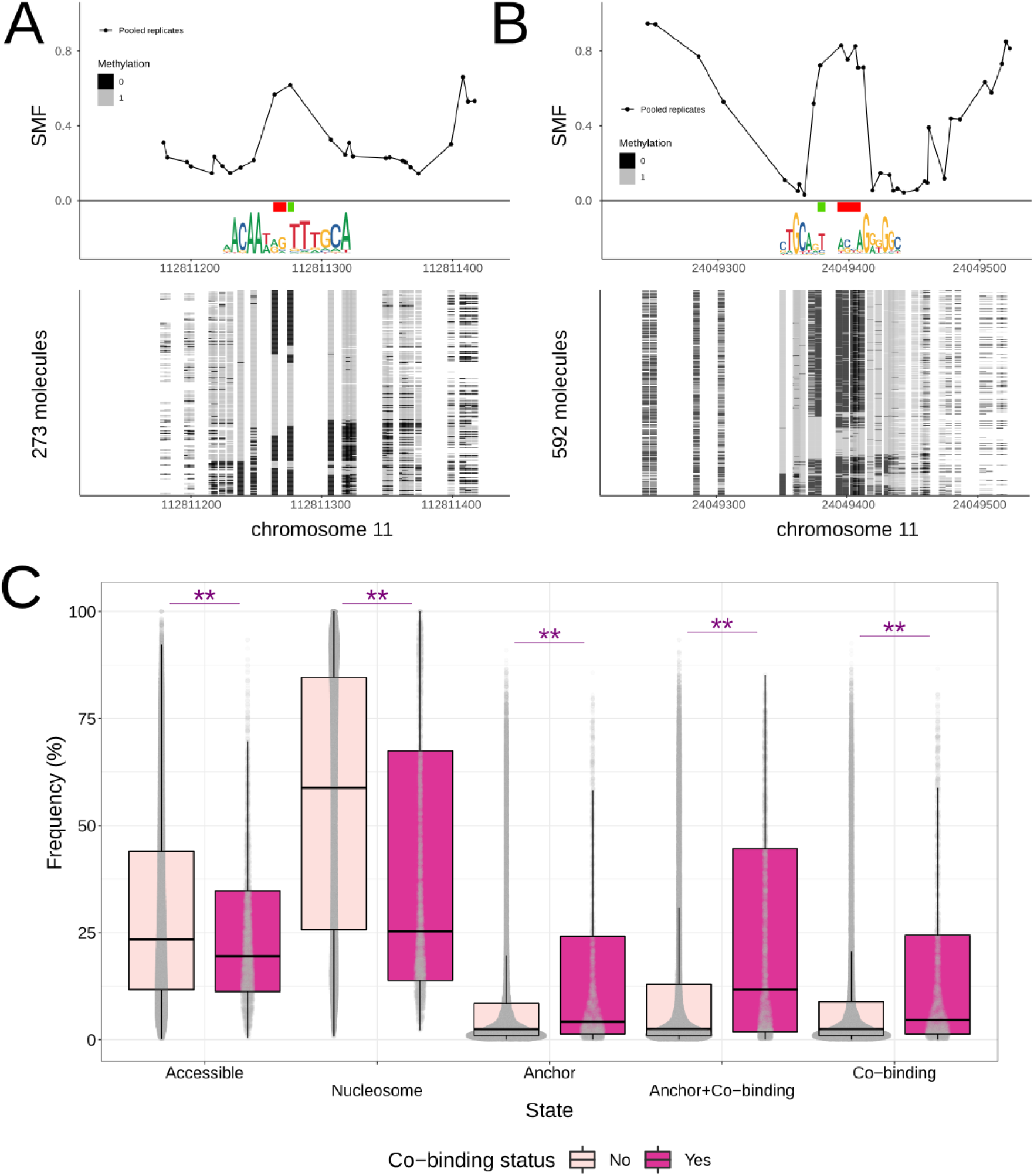
Single-molecule footprinting analysis. (**A-B**) Single-locus examples of predicted co-bound regions where the anchor TFBSs associate with SOX2 (A) and CTCF (B). Line plots in the top panels represent the average methylation levels. We provide logos and locations of the anchor motifs (red) and the predicted co-binding motifs (green). Stacks of sorted single molecules are visualized in the lower panels (light gray indicates methylated Cs, accessible positions; black - unmethylated Cs, protected). The y-axis corresponds to the total number of molecules. (**C**) We provide boxplots and dotplots showing the distribution of state frequencies of molecules that overlap the regions in two groups: one with predicted co-binding (shown in dark pink) and one with only an anchor TFBS (shown in light pink). The regions with co-binding events show a significantly higher frequency of molecules as co-occupied (“Anchor + Co-binding”). The differences between groups persist for molecules in other states. ** represents a Wilcoxon test p-value < 0.001.

## DISCUSSION

We introduced COBIND, a computational framework for discovering *de novo* DNA motif patterns in the vicinity of predetermined genomic regions. We applied COBIND to sets of TFBSs from UniBind to identify locally enriched DNA motifs that define co-binding patterns of cooperative TFs. The method uncovered both established and new co-binding patterns, with most TFs sharing a co-binding motif with other TFs from the same family. Additionally, we inferred, when possible, the TFs most likely co-binding the discovered patterns. We make the collection of co-binding patterns revealed by COBIND available to the scientific community for each of the 218 TFs and 57 TF families across seven species. The co-binding events captured by COBIND are likely functionally relevant since they exhibit higher evolutionary conservation than isolated TFBSs. Furthermore, chromatin and single-molecule footprinting data support the occurrence of the identified co-binding events on the same DNA molecules.

We used a combination of motif similarity and prior knowledge of PPIs to identify TFs that may cooperatively bind to the patterns discovered by COBIND. Identifying well-known TF co-binding events, such as POU5F1 with SOX2 or SOX17, essential pluripotency regulators (15, 16, 66), confirmed the approach’s efficacy. Importantly, we used a multi-species collection of DNA binding motifs to predict new co-binding configurations for pairs of TFs currently unsupported by PPIs. We discovered that NF-Y proteins are potential partners for HY5, ABF1, 3-4, and GBF2-3 TFs. Previous studies identified NF-Y proteins as co-binders for bZIP67 and bZIP28 proteins belonging to the same TF family (59–61). More specifically, other studies also pointed to physical interactions of either HY5 or ABF1, 3-4 with NF-YC9 proteins (67, 68). We identified the NF-Y-bound co-binding motifs using motifs associated with human TFs because DNA binding profiles for NF-Y are limited in *A. thaliana*. As HY5 competes with bZIP28, we acknowledge that the NF-Y binding pattern observed close to HY5 TFBSs might be occupied only when bZIP28 is binding. Nevertheless, using multi-species libraries generated potential co-binding predictions that will require further validations to decipher the binding cooperation of TFs at the corresponding genomic regions.

We observed that the COBIND predictions associated with CTCF anchor TFBSs exhibited a binding pattern that did not result from the co-binding of two TFs. Previous studies have discussed this binding pattern and have suggested that CTCF co-binds with ZBTB3 in mouse liver and various human cell lines, including embryonic stem cells and K562 (69, 70). Based on motif similarity, we inferred that ZBTB3 could also bind the additional motif identified by COBIND. Instead, the co-binding pattern identified by COBIND corresponds to an extended binding motif for CTCF, involving zinc fingers 9-11 (62). This orthogonal evidence allowed the two variants of extended CTCF motifs to complement the JASPAR database (MA1930.1 and MA1929.1). Furthermore, this binding pattern is critical for optimal CTCF protein residence time on DNA (62). Importantly, we found that the genomic regions harboring the extended motif were conserved in both mouse and human genomes, supporting the functional relevance of this binding pattern. Studies frequently reported that zinc-finger proteins bind DNA through a subset of their zinc fingers, but it remains unclear whether the other fingers have additional binding preferences (71). Thus, binding to extended motifs with additional zinc fingers may be a common characteristic of zinc finger proteins that *de novo* motif prediction tools like COBIND could capture.

The conservation of TFBSs is an essential indicator of their functional relevance in gene regulation (48). Furthermore, the combinatorial binding of TFs is important for evolutionary stability, and increased co-binding of TFs associates with a higher probability of a regulatory region being conserved (72). This study analyzed the conservation of co-binding patterns across different species. The regions predicted to harbor TFBSs with fixed spacing were more evolutionarily conserved than those with a single TFBS. Interestingly, not only were the predicted co-bound motif locations conserved, but the anchor motif locations also showed an increase in conservation. We found most co-binding patterns with significant conservation from the human and mouse datasets, which may reflect the more significant number of datasets analyzed for these species. Furthermore, there was a limited number of conservation tracks for other species, such as *A. thaliana,* where conservation was limited to flowering plants (49).

COBIND is a tool for the *de novo* discovery of co-occurring DNA patterns. It does not require *a priori* knowledge of DNA motifs, such as a known DNA motif binding library, to perform the discovery. The requirement of known motif collections as an input is a standard limitation for other tools dealing with TF binding analysis, such as SpaMo or TACO (20, 21). This reliance on existing collections of DNA binding motifs can be problematic as many TF binding motifs still need to be discovered or included. The only input required for COBIND is a set of genomic reference regions to “anchor” the analysis and generate regions where it will search for patterns. The *de novo* discovery can reveal potential new or unannotated DNA pattern motifs, thereby circumventing the constraints of other tools. As such, COBIND offers a powerful alternative to other motif discovery tools that rely on existing collections of DNA binding motifs. Nonetheless, a limitation is that additional analysis will be required to interpret newly discovered motifs.

In this study, we applied COBIND to genomic regions near TFBSs. However, one can use COBIND in different biological contexts. As a test case, we generated anchor regions for analysis by taking two nucleotides at the donor and acceptor sites of intron-exon boundaries in the human genome. Analysis with COBIND recovered known donor and acceptor motifs (Additional File 2: Figure S1), demonstrating the versatility of the approach for DNA pattern discovery in other types of biological problems.

We utilized the available SMF data to assess the co-occupancy of the anchor and co-binding motif instances on the same DNA molecules (53). A limitation of the assay is that one can only apply it to organisms and cell lines that can survive without endogenous methylation. We restricted our analyses to mouse ESCs, for which SMF was already available. A recent study assessed chromatin openness using methylation in the GpC context, allowing applications to different cell types with larger genomic coverage (73).

We recognize that COBIND and this study have several limitations. A fundamental limitation of COBIND is the restricted prediction of co-binding motifs with fixed spacing with the anchor motif. Because of the strict spacing requirements and the limitations of the NMF, COBIND restricts the search space to the close proximity of TFBSs. Previous studies have hypothesized the existence of two distinct sets of enhancers with distinct information-processing mechanisms (10). One proposed mechanism distinguishes between enhancers where TF binding is highly cooperative and coordinated with fixed spacing, referred to as the ’enhanceosome’ model, and enhancers where TF cooperation is flexible, known as the ’billboard’ model (74). Regardless, these mechanisms are compatible with the presence of both fixed and flexible cooperation of TFs at regulatory elements, depending on the type of TF cooperation. Recent studies support both mechanisms and even though some suggest that TFBS spacing and orientation are not key determinants of transcription regulation, other studies contradict this statement (10–14). The evolutionary conservation of the co-binding patterns revealed by COBIND further argues for the functional relevance of strict grammar.

COBIND relies on the NMF algorithm to discover the co-binding patterns, so it requires enough sequences harboring the fixed pattern. Our simulated data showed that we obtained reliable results when at least 3% of at least 1000 sequences contained the pattern to be predicted by COBIND. Since we considered the anchor TFBS and the flanks independently in the COBIND processing, discovering overlapping motifs represents a challenge. COBIND could detect such overlaps by revealing a small co-binding motif, but interpreting such results becomes increasingly difficult as the partial motif becomes small. Furthermore, we used anchor TFBSs predicted from TF binding profiles corresponding to the canonical motifs recognized by the corresponding TFs. This approach prohibits the identification of co-binding patterns where the anchor TFs would recognize altered motifs when cooperating with other proteins. Finally, another limitation lies in the computational time necessary for motif clustering when the NMF identifies many possible motifs. Our comparison to other tools revealed that COBIND’s computational time was not prohibitive.

## DATA AVAILABILITY

COBIND is implemented in Python, R, and C++. The COBIND source code and documentation are available at https://bitbucket.org/CBGR/cobind_tool/src/master/. The source code and data to reproduce the results outlined in this report is available at https://bitbucket.org/CBGR/cobind_manuscript/src/master/ and pre-processed data deposited at https://doi.org/10.5281/zenodo.7681483. All results presented here are available to the community through a webpage at https://cbgr.bitbucket.io/COBIND_2023/.

## AUTHORS’ CONTRIBUTIONS

We follow here the Contributor Roles Taxonomy (CRediT) (75). Ieva Rauluseviciute: methodology, software, validation, investigation, formal analysis, writing - original draft, visualization; Timothée Launay: methodology, investigation, software, writing - review & editing; Guido Barzaghi: resources, writing - review & editing; Sarvesh Nikumbh: writing - review & editing; Boris Lenhard: writing - review & editing, funding acquisition; Arnaud Regis Krebs: writing - review & editing, funding acquisition; Jaime A. Castro-Mondragon: conceptualization, methodology, writing - review & editing; Anthony Mathelier: conceptualization, methodology, writing - review & editing, supervision, project administration, funding acquisition.

## ACKNOWLEDGMENTS

As “research parasites” (76), we thank all the researchers who made their data available. We thank Marcel Schulz for his suggestion on protein-protein interaction data analysis, François Parcy for fruitful discussion on the plant co-binding motifs and his help finding the plant evolutionary conservation data, Judith Zaugg for providing feedback on the project, Roza Berhanu Lemma, Vipin Kumar, Ladislav Hovan, and Rafael Riudavets Puig for reading the manuscript and providing valuable feedback, Harold Gutch, Torfinn Nome, and the NCMM IT team for their IT support, Ingrid Kjelsvik for administrative support, and the members of the Kuijjer and Mathelier groups for insightful discussions.

## REFERENCES

1. Wingender, E., Schoeps, T., Haubrock, M., Krull, M. and Dönitz, J. (2018) TFClass: expanding the classification of human transcription factors to their mammalian orthologs. Nucleic Acids Res., 46, D343–D347.

2. Lambert, S.A., Jolma, A., Campitelli, L.F., Das, P.K., Yin, Y., Albu, M., Chen, X., Taipale, J., Hughes, T.R. and Weirauch, M.T. (02/2018) The Human Transcription Factors. Cell, 172, 650–665.

3. Zeitlinger, J. (2020) Seven myths of how transcription factors read the cis-regulatory code. Current Opinion in Systems Biology, 23, 22–31.

4. Suter, D.M. (2020) Transcription Factors and DNA Play Hide and Seek. Trends Cell Biol., 30, 491–500.

5. Weirauch, M.T., Yang, A., Albu, M., Cote, A.G., Montenegro-Montero, A., Drewe, P., Najafabadi, H.S., Lambert, S.A., Mann, I., Cook, K., et al. (2014) Determination and inference of eukaryotic transcription factor sequence specificity. Cell, 158, 1431–1443.

6. Kulakovskiy, I.V., Vorontsov, I.E., Yevshin, I.S., Sharipov, R.N., Fedorova, A.D., Rumynskiy, E.I., Medvedeva, Y.A., Magana-Mora, A., Bajic, V.B., Papatsenko, D.A., et al. (2018) HOCOMOCO: towards a complete collection of transcription factor binding models for human and mouse via large-scale ChIP-Seq analysis. Nucleic Acids Res., 46, D252–D259.

7. Castro-Mondragon, J.A., Riudavets-Puig, R., Rauluseviciute, I., Berhanu Lemma, R., Turchi, L., Blanc-Mathieu, R., Lucas, J., Boddie, P., Khan, A., Manosalva Pérez, N., et al. (2021) JASPAR 2022: the 9th release of the open-access database of transcription factor binding profiles. Nucleic Acids Res.

8. Reiter, F., Wienerroither, S. and Stark, A. (2017) Combinatorial function of transcription factors and cofactors. Curr. Opin. Genet. Dev., 43, 73–81.

9. Zhou, M., Li, H., Wang, X. and Guan, Y. (2020) Evidence of widespread, independent sequence signature for transcription factor cobinding. Genome Res.

10. Arnosti, D.N. and Kulkarni, M.M. (2005) Transcriptional enhancers: Intelligent enhanceosomes or flexible billboards? J. Cell. Biochem., 94, 890–898.

11. King, D.M., Hong, C.K.Y., Shepherdson, J.L., Granas, D.M., Maricque, B.B. and Cohen, B.A. (2020) Synthetic and genomic regulatory elements reveal aspects of cis-regulatory grammar in mouse embryonic stem cells. Elife, 9, e41279.

12. Avsec, Ž., Weilert, M., Shrikumar, A., Krueger, S., Alexandari, A., Dalal, K., Fropf, R., McAnany, C., Gagneur, J., Kundaje, A., et al. (2021) Base-resolution models of transcription-factor binding reveal soft motif syntax. Nat. Genet., 53, 354–366.

13. Sahu, B., Hartonen, T., Pihlajamaa, P., Wei, B., Dave, K., Zhu, F., Kaasinen, E., Lidschreiber, K., Lidschreiber, M., Daub, C.O., et al. (2022) Sequence determinants of human gene regulatory elements. Nat. Genet., 54, 283–294.

14. Georgakopoulos-Soares, I., Deng, C., Agarwal, V., Chan, C.S.Y., Zhao, J., Inoue, F. and Ahituv, N. (2023) Transcription factor binding site orientation and order are major drivers of gene regulatory activity. Nat. Commun., 14, 2333.

15. Li, M. and Belmonte, J.C.I. (2018-4) Deconstructing the pluripotency gene regulatory network. Nat. Cell Biol., 20, 382– 392.

16. Aksoy, I., Jauch, R., Chen, J., Dyla, M., Divakar, U., Bogu, G.K., Teo, R., Leng Ng, C.K., Herath, W., Lili, S., et al. (2013) Oct4 switches partnering from Sox2 to Sox17 to reinterpret the enhancer code and specify endoderm. EMBO J., 32, 938–953.

17. Nagy, G. and Nagy, L. (2020) Motif grammar: The basis of the language of gene expression. Comput. Struct. Biotechnol. J., 18, 2026–2032.

18. Jauch, R., Aksoy, I., Hutchins, A.P., Ng, C.K.L., Tian, X.F., Chen, J., Palasingam, P., Robson, P., Stanton, L.W. and Kolatkar, P.R. (2011) Conversion of Sox17 into a pluripotency reprogramming factor by reengineering its association with Oct4 on DNA. Stem Cells, 29, 940–951.

19. Jolma, A., Yin, Y., Nitta, K.R., Dave, K., Popov, A., Taipale, M., Enge, M., Kivioja, T., Morgunova, E. and Taipale, J. (2016) DNA-dependent formation of transcription factor pairs alters their binding specificity. Nature, 534, S15–S16.

20. Jankowski, A., Prabhakar, S. and Tiuryn, J. (2014) TACO: a general-purpose tool for predicting cell-type-specific transcription factor dimers. BMC Genomics, 15, 208.

21. Whitington, T., Frith, M.C., Johnson, J. and Bailey, T.L. (2011-8) Inferring transcription factor complexes from ChIP-seq data. Nucleic Acids Res., 39, e98.

21. Levitsky, V., Zemlyanskaya, E., Oshchepkov, D., Podkolodnaya, O., Ignatieva, E., Grosse, I., Mironova, V. and Merkulova, T. (2019) A single ChIP-seq dataset is sufficient for comprehensive analysis of motifs co-occurrence with MCOT package. Nucleic Acids Res., 47, e139.

23. Bentsen, M., Heger, V., Schultheis, H., Kuenne, C. and Looso, M. (2022) TF-COMB - Discovering grammar of transcription factor binding sites. Comput. Struct. Biotechnol. J., 20, 4040–4051.

24. Park, P.J. (10/2009) ChIP–seq: advantages and challenges of a maturing technology. Nat. Rev. Genet., 10, 669–680.

25. van Helden, J., Rios, A.F. and Collado-Vides, J. (2000) Discovering regulatory elements in non-coding sequences by analysis of spaced dyads. Nucleic Acids Res., 28, 1808–1818.

26. Defrance, M., Janky, R. ’s, Sand, O. and van Helden, J. (2008) Using RSAT oligo-analysis and dyad-analysis tools to discover regulatory signals in nucleic sequences. Nat. Protoc., 3, 1589–1603.

27. Minnoye, L., Taskiran, I.I., Mauduit, D., Fazio, M., Van Aerschot, L., Hulselmans, G., Christiaens, V., Makhzami, S., Seltenhammer, M., Karras, P., et al. (2020) Cross-species analysis of enhancer logic using deep learning. Genome Res., 30, 1815–1834.

28. Stein-O’Brien, G.L., Arora, R., Culhane, A.C., Favorov, A.V., Garmire, L.X., Greene, C.S., Goff, L.A., Li, Y., Ngom, A., Ochs, M.F., et al. (2018) Enter the Matrix: Factorization Uncovers Knowledge from Omics. Trends Genet., 34, 790– 805.

29. Lee, D.D. and Seung, H.S. (1999) Learning the parts of objects by non-negative matrix factorization. Nature, 401, 788– 791.

30. Devarajan, K. (2008) Nonnegative Matrix Factorization: An Analytical and Interpretive Tool in Computational Biology. PLoS Comput. Biol., 4, e1000029.

31. Nikumbh, S. and Lenhard, B. (2023) Identifying promoter sequence architectures via a chunking-based algorithm using non-negative matrix factorisation. bioRxiv, 10.1101/2023.03.02.530868.

32. Mölder, F., Jablonski, K.P., Letcher, B., Hall, M.B., Tomkins-Tinch, C.H., Sochat, V., Forster, J., Lee, S., Twardziok, S.O., Kanitz, A., et al. (2021) Sustainable data analysis with Snakemake. F1000Res., 10, 33.

33. Puig, R.R., Boddie, P., Khan, A., Castro-Mondragon, J.A. and Mathelier, A. (2021) UniBind: maps of high-confidence direct TF-DNA interactions across nine species. BMC Genomics, 22, 482.

34. Quinlan, A.R. and Hall, I.M. (2010) BEDTools: a flexible suite of utilities for comparing genomic features. Bioinformatics, 26, 841–842.

35. Pedregosa, F. (2011) Scikit-learn: Machine Learning in Python. J. Mach. Learn. Res., 12, 2825–2830.

36. Kim, H. and Park, H. (2007) Sparse non-negative matrix factorizations via alternating non-negativity-constrained least squares for microarray data analysis. Bioinformatics, 23, 1495–1502.

37. Gini, C. (1912) Variabilità e mutabilità: contributo allo studio delle distribuzioni e delle relazioni statistiche. [Fasc. I.] Tipogr. di P. Cuppini.

38. Castro-Mondragon, J.A., Jaeger, S., Thieffry, D., Thomas-Chollier, M. and van Helden, J. (2017) RSAT matrix-clustering: dynamic exploration and redundancy reduction of transcription factor binding motif collections. Nucleic Acids Res., 45, e119.

39. Castro-Mondragon, J. (2022) matrix-clustering_stand-alone.

40. Shrikumar, A., Tian, K., Avsec, Ž., Shcherbina, A., Banerjee, A., Sharmin, M., Nair, S. and Kundaje, A. (2020) Technical Note on Transcription Factor Motif Discovery from Importance Scores (TF-MoDISco) version 0.5.6.5.

41. Gupta, S., Stamatoyannopoulos, J.A., Bailey, T.L. and Noble, W.S. (2007) Quantifying similarity between motifs. Genome Biol., 8, R24.

42. Khan, A., Riudavets Puig, R., Boddie, P. and Mathelier, A. (2020) BiasAway: command-line and web server to generate nucleotide composition-matched DNA background sequences. Bioinformatics, 10.1093/bioinformatics/btaa928.

43. Machanick, P. and Bailey, T.L. (2011) MEME-ChIP: motif analysis of large DNA datasets. Bioinformatics, 27, 1696–1697.

44. Thomas-Chollier, M., Darbo, E., Herrmann, C., Defrance, M., Thieffry, D. and van Helden, J. (2012) A complete workflow for the analysis of full-size ChIP-seq (and similar) data sets using peak-motifs. Nat. Protoc., 7, 1551–1568.

45. Thomas-Chollier, M., Herrmann, C., Defrance, M., Sand, O., Thieffry, D. and van Helden, J. (2012) RSAT peak-motifs: motif analysis in full-size ChIP-seq datasets. Nucleic Acids Res., 40, e31–e31.

46. Fornes, O., Castro-Mondragon, J.A., Khan, A., van der Lee, R., Zhang, X., Richmond, P.A., Modi, B.P., Correard, S., Gheorghe, M., Baranašić, D., et al. (2020) JASPAR 2020: update of the open-access database of transcription factor binding profiles. Nucleic Acids Res., 48, D87–D92.

47. Szklarczyk, D., Gable, A.L., Lyon, D., Junge, A., Wyder, S., Huerta-Cepas, J., Simonovic, M., Doncheva, N.T., Morris, J.H., Bork, P., et al. (2019) STRING v11: protein-protein association networks with increased coverage, supporting functional discovery in genome-wide experimental datasets. Nucleic Acids Res., 47, D607–D613.

48. Siepel, A., Bejerano, G., Pedersen, J.S., Hinrichs, A.S., Hou, M., Rosenbloom, K., Clawson, H., Spieth, J., Hillier, L.W., Richards, S., et al. (2005) Evolutionarily conserved elements in vertebrate, insect, worm, and yeast genomes. Genome Res., 15, 1034–1050.

49. Tian, F., Yang, D.-C., Meng, Y.-Q., Jin, J. and Gao, G. (2020) PlantRegMap: charting functional regulatory maps in plants. Nucleic Acids Res., 48, D1104–D1113.

50. Pohl, A. and Beato, M. (2014) bwtool: a tool for bigWig files. Bioinformatics, 30, 1618–1619.

51. Vierstra, J., Lazar, J., Sandstrom, R., Halow, J., Lee, K., Bates, D., Diegel, M., Dunn, D., Neri, F., Haugen, E., et al. (2020) Global reference mapping of human transcription factor footprints. Nature, 583, 729–736.

52. Sönmezer, C., Kleinendorst, R., Imanci, D., Barzaghi, G., Villacorta, L., Schübeler, D., Benes, V., Molina, N. and Krebs, A.R. (2021) Molecular Co-occupancy Identifies Transcription Factor Binding Cooperativity In Vivo. Mol. Cell, 81, 255– 267.e6.

53. Kleinendorst, R.W.D., Barzaghi, G., Smith, M.L., Zaugg, J.B. and Krebs, A.R. (2021) Genome-wide quantification of transcription factor binding at single-DNA-molecule resolution using methyl-transferase footprinting. Nat. Protoc., 16, 5673–5706.

54. Barzaghi, G., Krebs, A. and Smith, M. (2022) SingleMoleculeFootprinting: Analysis tools for Single Molecule Footprinting (SMF) data Bioconductor version: Release (3.15).

55. Mistri, T.K., Devasia, A.G., Chu, L.T., Ng, W.P., Halbritter, F., Colby, D., Martynoga, B., Tomlinson, S.R., Chambers, I., Robson, P., et al. (2015) Selective influence of Sox2 on POU transcription factor binding in embryonic and neural stem cells. EMBO Rep., 16, 1177–1191.

56. Jiang, S.W., Desai, D., Khan, S. and Eberhardt, N.L. (2000) Cooperative binding of TEF-1 to repeated GGAATG-related consensus elements with restricted spatial separation and orientation. DNA Cell Biol., 19, 507–514.

57. Anbanandam, A., Albarado, D.C., Nguyen, C.T., Halder, G., Gao, X. and Veeraraghavan, S. (2006) Insights into transcription enhancer factor 1 (TEF-1) activity from the solution structure of the TEA domain. Proc. Natl. Acad. Sci. U. S. A., 103, 17225–17230.

58. Lee, D.-S., Vonrhein, C., Albarado, D., Raman, C.S. and Veeraraghavan, S. (2016) A potential structural switch for regulating DNA-binding by TEAD transcription factors. J. Mol. Biol., 428, 2557–2568.

59. Mendes, A., Kelly, A.A., van Erp, H., Shaw, E., Powers, S.J., Kurup, S. and Eastmond, P.J. (2013) bZIP67 Regulates the Omega-3 Fatty Acid Content of Arabidopsis Seed Oil by Activating FATTY ACID DESATURASE3. Plant Cell, 25, 3104–3116.

60. Pastor-Cantizano, N., Ko, D.K., Angelos, E., Pu, Y. and Brandizzi, F. (2020-2) Functional diversification of ER stress responses in Arabidopsis. Trends Biochem. Sci., 45, 123–136.

61. Nawkar, G.M., Kang, C.H., Maibam, P., Park, J.H., Jung, Y.J., Chae, H.B., Chi, Y.H., Jung, I.J., Kim, W.Y., Yun, D.-J., et al. (2017) HY5, a positive regulator of light signaling, negatively controls the unfolded protein response in Arabidopsis. Proc. Natl. Acad. Sci. U. S. A., 114, 2084–2089.

62. Soochit, W., Sleutels, F., Stik, G., Bartkuhn, M., Basu, S., Hernandez, S.C., Merzouk, S., Vidal, E., Boers, R., Boers, J., et al. (2021) CTCF chromatin residence time controls three-dimensional genome organization, gene expression and DNA methylation in pluripotent cells. Nat. Cell Biol., 23, 881–893.

63. Nakahashi, H., Kieffer Kwon, K.-R., Resch, W., Vian, L., Dose, M., Stavreva, D., Hakim, O., Pruett, N., Nelson, S., Yamane, A., et al. (2013) A genome-wide map of CTCF multivalency redefines the CTCF code. Cell Rep., 3, 1678– 1689.

64. Thurman, R.E., Rynes, E., Humbert, R., Vierstra, J., Maurano, M.T., Haugen, E., Sheffield, N.C., Stergachis, A.B., Wang, H., Vernot, B., et al. (2012) The accessible chromatin landscape of the human genome. Nature, 489, 75–82.

65. Funk, C.C., Casella, A.M., Jung, S., Richards, M.A., Rodriguez, A., Shannon, P., Donovan-Maiye, R., Heavner, B., Chard, K., Xiao, Y., et al. (2020) Atlas of Transcription Factor Binding Sites from ENCODE DNase Hypersensitivity Data across 27 Tissue Types. Cell Rep., 32, 108029.

66. Pan, X., Cang, X., Dan, S., Li, J., Cheng, J., Kang, B., Duan, X., Shen, B. and Wang, Y.-J. (2016) Site-specific Disruption of the Oct4/Sox2 Protein Interaction Reveals Coordinated Mesendodermal Differentiation and the Epithelial-Mesenchymal Transition. J. Biol. Chem., 291, 18353–18369.

67. Kumimoto, R.W., Siriwardana, C.L., Gayler, K.K., Risinger, J.R., Siefers, N. and Holt, B.F., 3rd (2013) NUCLEAR FACTOR Y transcription factors have both opposing and additive roles in ABA-mediated seed germination. PLoS One, 8, e59481.

68. Myers, Z.A., Kumimoto, R.W., Siriwardana, C.L., Gayler, K.K., Risinger, J.R., Pezzetta, D. and Holt, B.F., Iii (2016) NUCLEAR FACTOR Y, Subunit C (NF-YC) Transcription Factors Are Positive Regulators of Photomorphogenesis in Arabidopsis thaliana. PLoS Genet., 12, e1006333.

69. Wang, A.W., Wang, Y.J., Zahm, A.M., Morgan, A.R., Wangensteen, K.J. and Kaestner, K.H. (2020) The Dynamic Chromatin Architecture of the Regenerating Liver. Cellular and Molecular Gastroenterology and Hepatology, 9, 121– 143.

70. Boyle, A.P., Song, L., Lee, B.-K., London, D., Keefe, D., Birney, E., Iyer, V.R., Crawford, G.E. and Furey, T.S. (2011) High-resolution genome-wide in vivo footprinting of diverse transcription factors in human cells. Genome Res., 21, 456–464.

71. Schmitges, F.W., Radovani, E., Najafabadi, H.S., Barazandeh, M., Campitelli, L.F., Yin, Y., Jolma, A., Zhong, G., Guo, H., Kanagalingam, T., et al. (2016) Multiparameter functional diversity of human C2H2 zinc finger proteins. Genome Res., 26, 1742–1752.

72. Stefflova, K., Thybert, D., Wilson, M.D., Streeter, I., Aleksic, J., Karagianni, P., Brazma, A., Adams, D.J., Talianidis, I., Marioni, J.C., et al. (2013) Cooperativity and rapid evolution of cobound transcription factors in closely related mammals. Cell, 154, 530–540.

73. Kreibich, E., Kleinendorst, R., Barzaghi, G., Kaspar, S. and Krebs, A.R. (2023) Single-molecule footprinting identifies context-dependent regulation of enhancers by DNA methylation. Mol. Cell, 83, 787–802.e9.

74. Slattery, M., Zhou, T., Yang, L., Dantas Machado, A.C., Gordân, R. and Rohs, R. (09/2014) Absence of a simple code: how transcription factors read the genome. Trends Biochem. Sci., 39, 381–399.

75. Brand, A., Allen, L., Altman, M., Hlava, M. and Scott, J. (2015) Beyond authorship: attribution, contribution, collaboration, and credit. Learn. Publ., 28, 151–155.

76. Longo, D.L. and Drazen, J.M. (2016) Data Sharing. N. Engl. J. Med., 374, 276–277.

