## Additional File 1 for "Identification of transcription factor co-binding patterns with non-negative matrix factorization"

### Supplementary figures

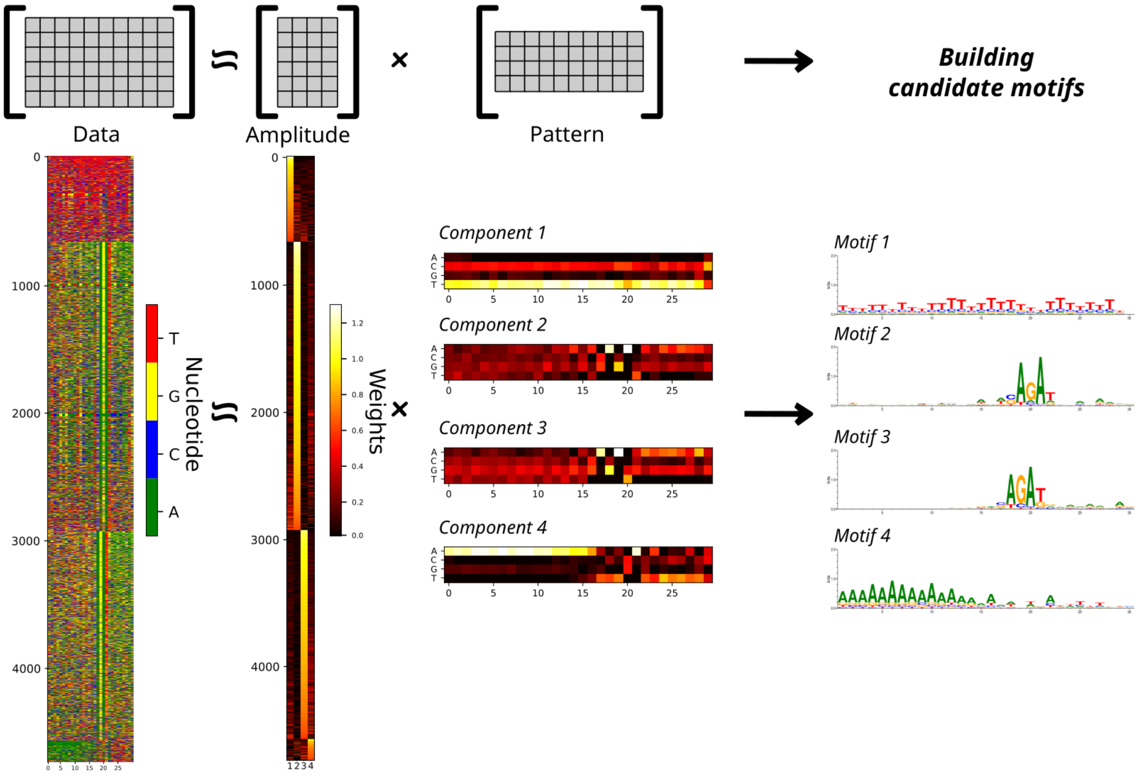


***Figure S1*. Non-negative matrix factorization discovers sequence patterns from one-hot encoded sequences.** The factorization results in amplitude and pattern matrices with a defined number of components *k* (in this example, *k=4*). The pattern matrices reveal candidate pattern motifs. We seek motifs with highly condensed information content (IC) surrounded by positions with low IC. For example, motifs 2 and 3 have a higher Gini coefficient than motifs 1 and 4, for which we observe an even distribution over IC the entire motif length.

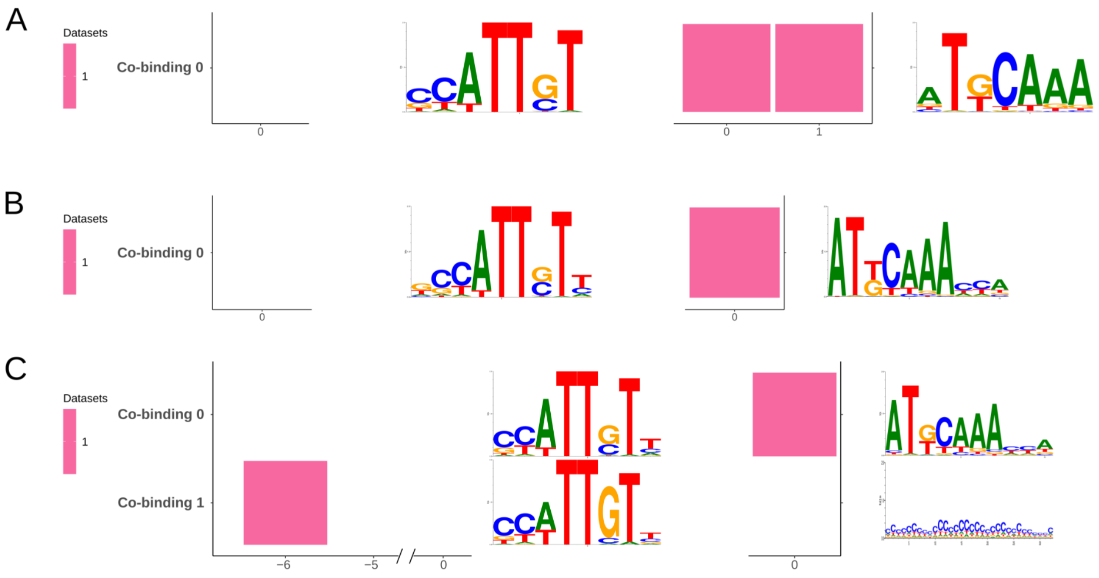


***Figure S2*. Investigating larger flanking regions results in noisier co-binding motifs.** The figure presents the co-binding patterns discovered by COBIND when applied to genomic regions flanking the anchor motif, considering 30bp (**A**), 50bp (**B**), and 100bp (**C**). Considering 50bp captures only one spacing configuration. Considering 100bp discovered an extra non-informative motif.

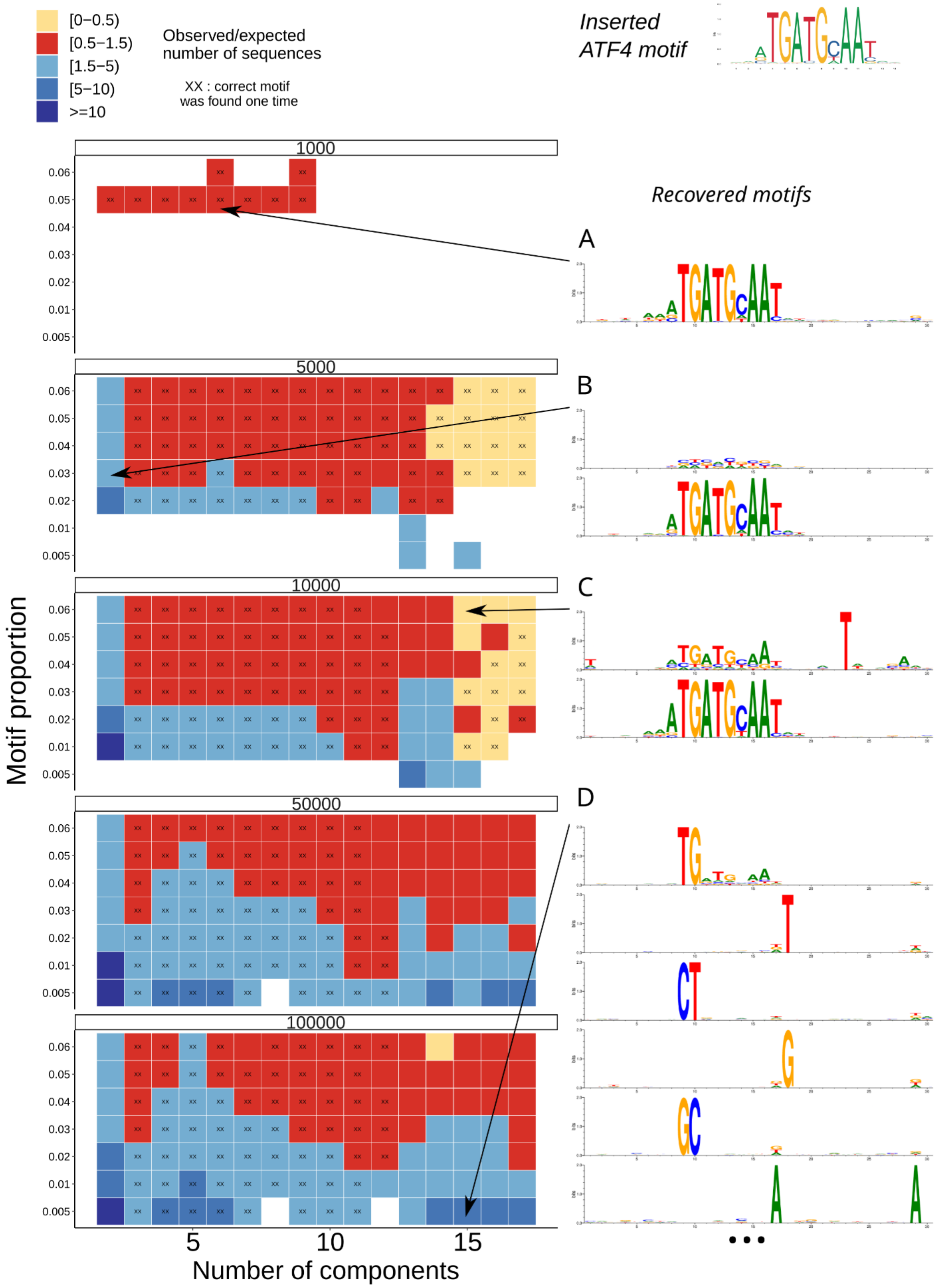

[0–0.5)
[0.5–1.5)
[1.5–5)
[5–10)
>=10
Observed/expected number of sequences
XX : correct motif was found one time
Inserted ATF4 motif
Recovered motifs
A
B
C
D
1000
5000
10000
50000
100000
Motif proportion
Number of components

***Figure S3*. Simulation experiments.** We used simulated random sequences to introduce instances of a known motif (JASPAR MA0833.2 TF binding profile) to run NMF (see Methods). (**A**) The inserted motif is captured in many instances, and the ratio of observed over expected instances is within [0.5, 1.5]. (**B**) When considering two components when applying the NMF, more sequences are associated with the motif, which makes the motif noisier. (**C**) With larger numbers of components, the recovered motifs derive from fewer sequences than the number of sequences containing the motif, resulting in a motif discovered multiple times. (**D**) With a sufficiently large number of components, the injected motif gets decomposed over numerous clusters, which leads to partial motif discovery.

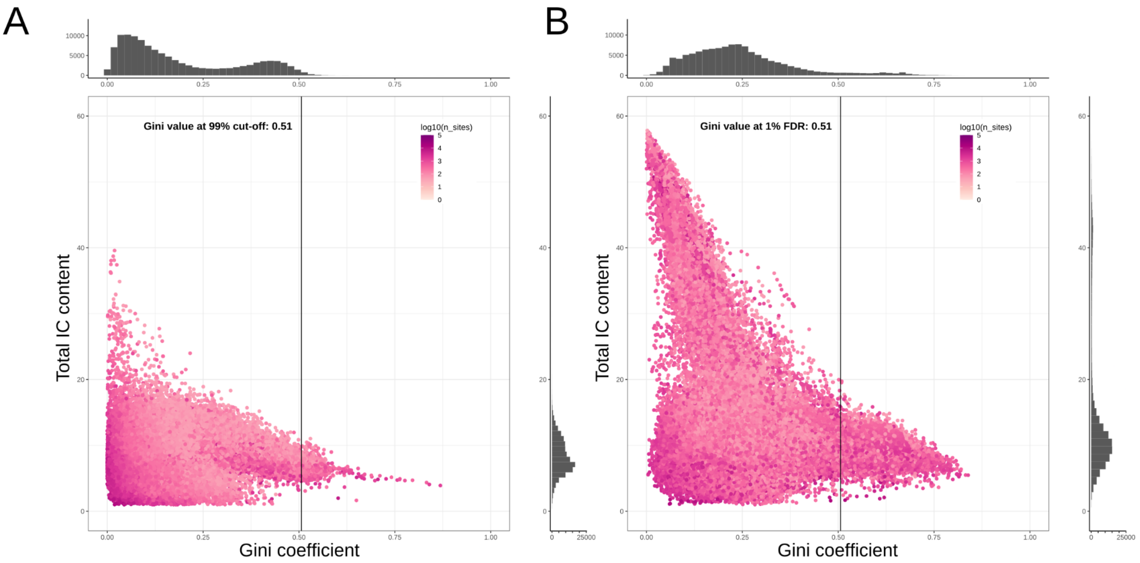


***Figure S4.*** **Species-specific Gini coefficient thresholding using simulated human data.** (**A**) Distribution of Gini coefficients and information content (IC) values for the motifs discovered in simulated data (see Methods). The vertical line indicates the 99th percentile of the Gini distribution (equal to 0.5). (**B**) Distribution of Gini coefficients and information content (IC) values for the motifs discovered in the real human data. The vertical line represents a Gini coefficient cut-off value of 0.5, derived from the simulated data (A).

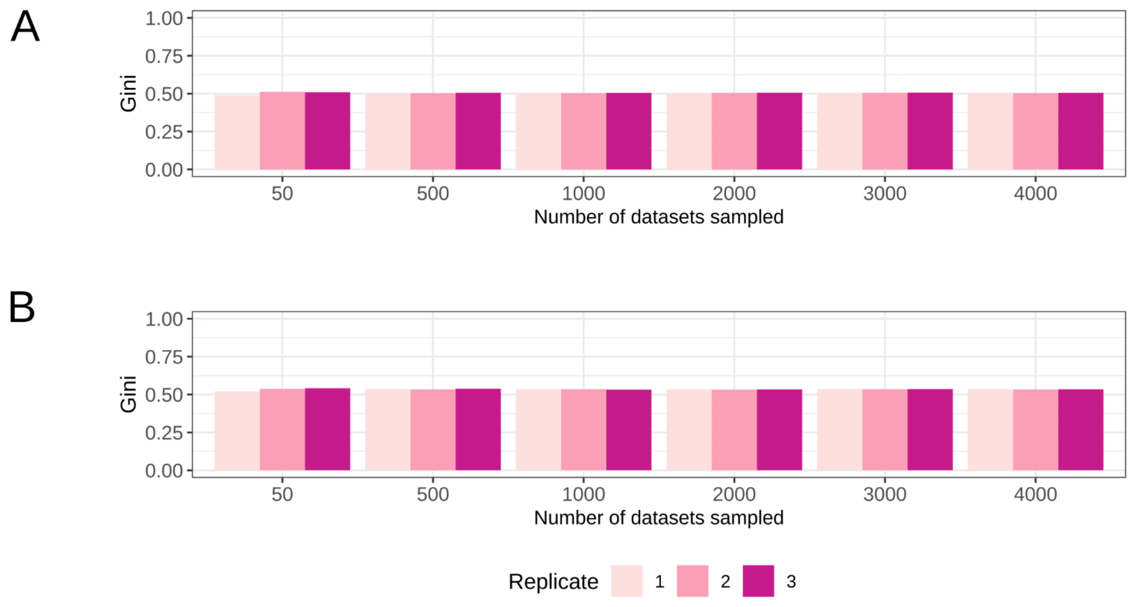


***Figure S5*. Consistency of the Gini coefficient thresholds.** We observed consistent Gini coefficient thresholds (y-axis) derived from human (**A**) and mouse (**B**) simulated data from different numbers of datasets (x-axis) with three replicates.

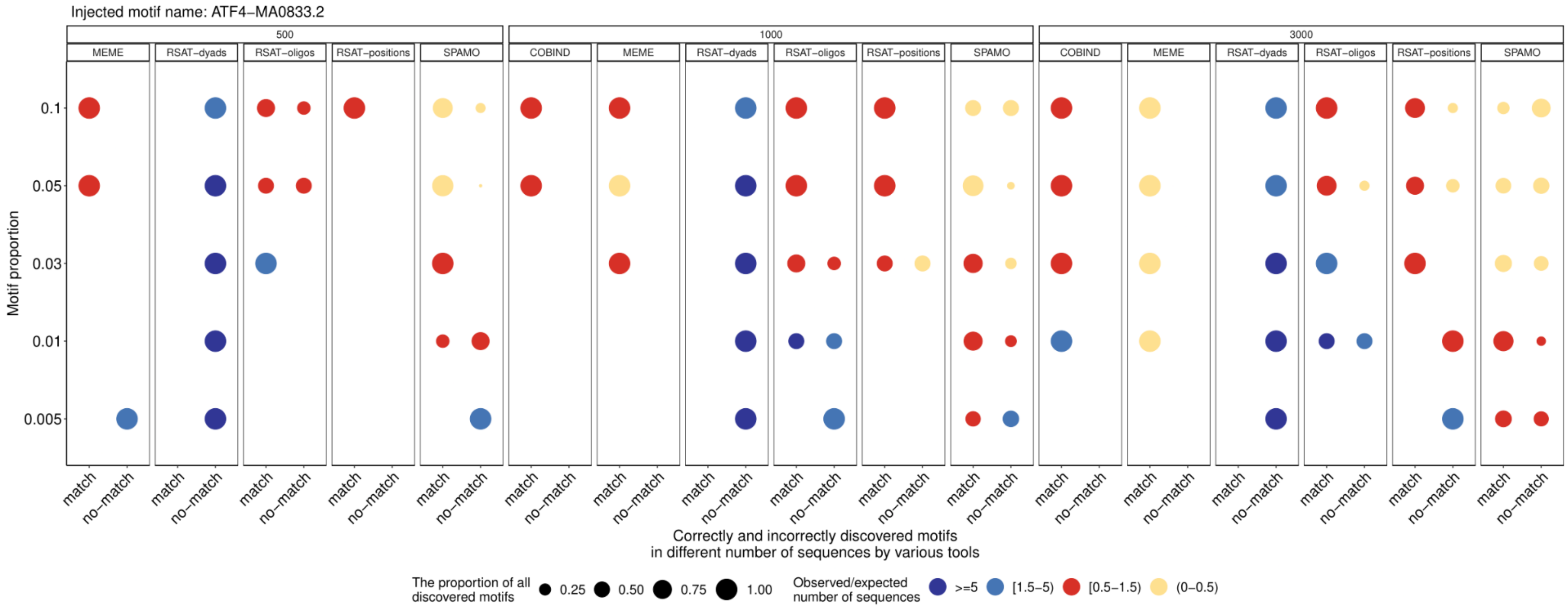


*(Continued on the next page)*

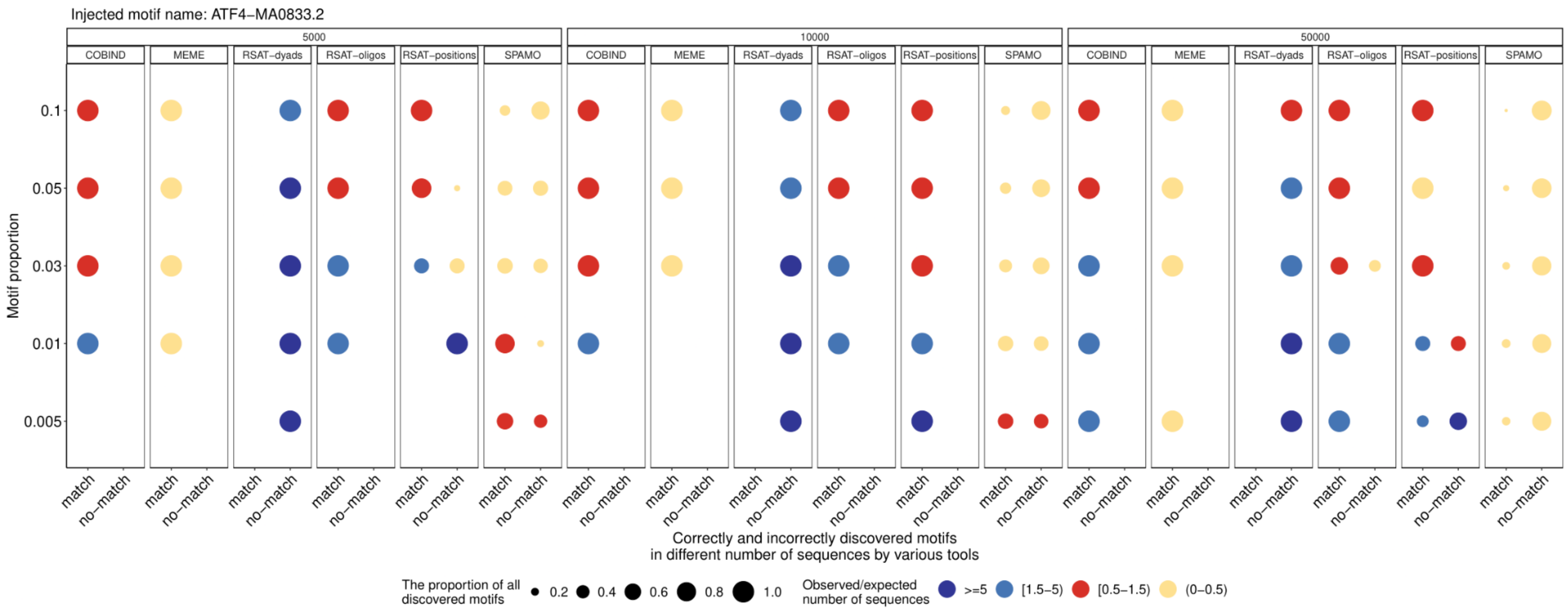


***Figure S6.*** **Comparisons between COBIND and other tools on simulated data.** We ran COBIND, MEME, RSAT-dyads, RSAT-oligos, RSAT-positions, and SPAMO (subfacets) on simulated data containing different numbers of sequences (from 500 to 50,000, see facets) (see Methods for details). We considered (i) if the discovered motifs were similar to the original inserted one ("match", "no-match", see Methods), (ii) the ratio between the number of predicted sequences containing the predicted motif (observed) and the expected number of sequences with the motif (rows; see color coding for "observed/expected" number of sequences with a motif), and (iii) the proportion of "match" motifs in all discovered motifs for each dataset. A good performance when there are only "match" motifs discovered or the proportion of "match" motifs is bigger (larger points) than "no-match" motifs, and the observed/expected ratio is in the 0.5-1.5 range (red colored points). Note that no motif was predicted by COBIND when using 500 sequences.

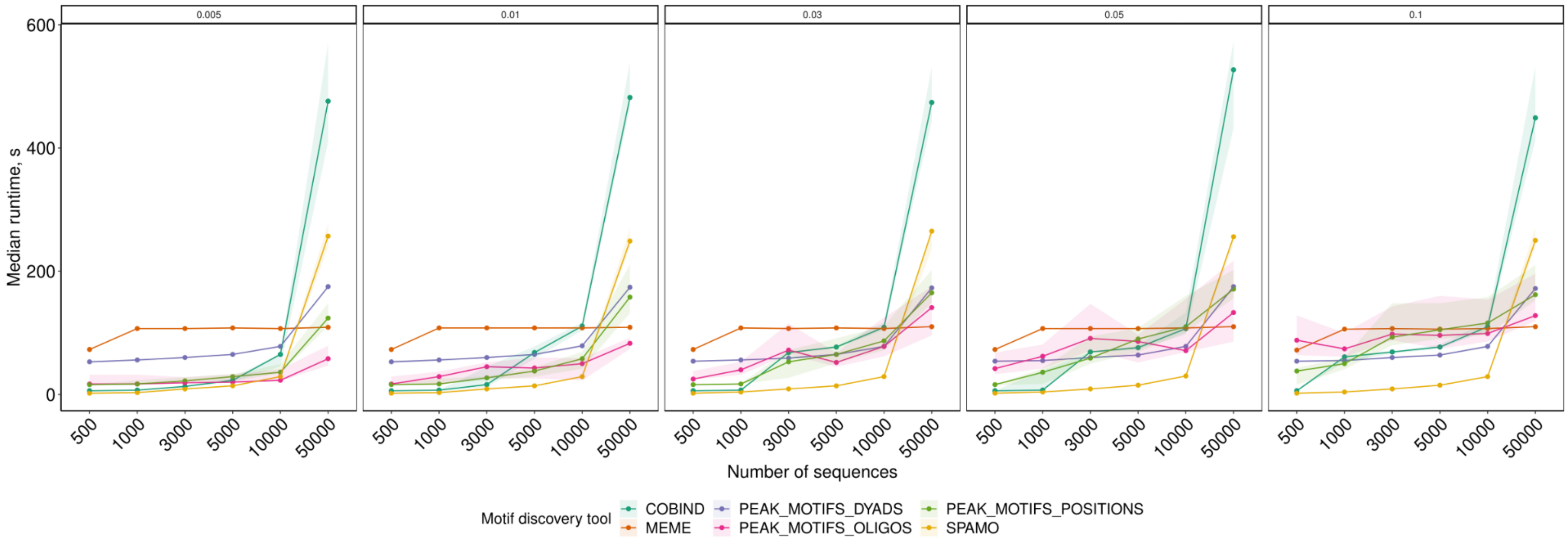


***Figure S7.*** **Run time comparison on simulated data.** The figure reports the running time (y-axis) of the different tools (see legend) when applied to different numbers of sequences (columns) with distinct proportions of injected motif instances (see facets).

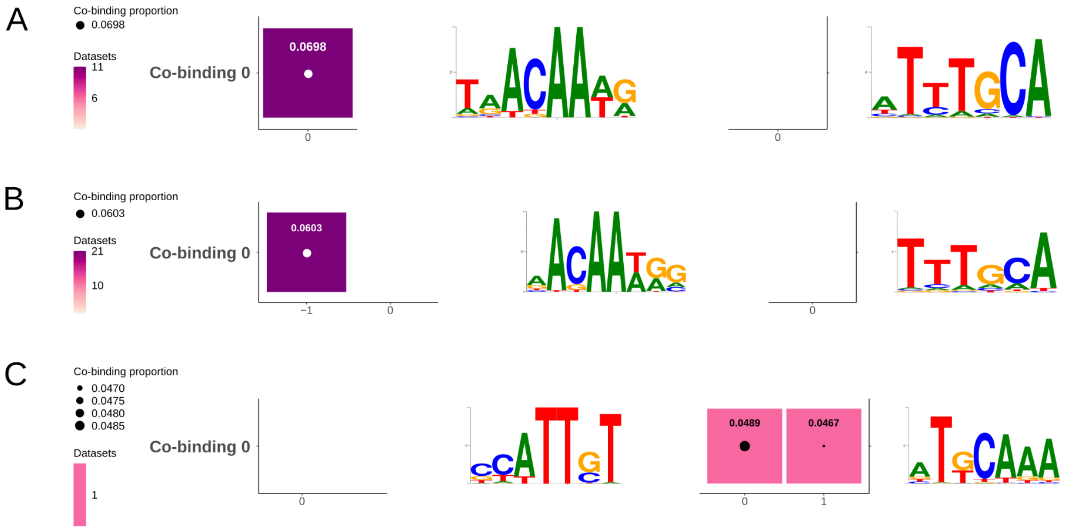


***Figure S8.*** **Applying COBIND to SOX2 and SOX17 *Homo sapiens* and *Mus musculus* TFBS datasets.** POU5F1 partners with SOX2 to bind its canonical motif in *H. sapiens* (**A**) and *M. musculus* (**B**). In contrast, POU5F1 associates with SOX17 to bind a compressed motif in *M. musculus* (**C**).

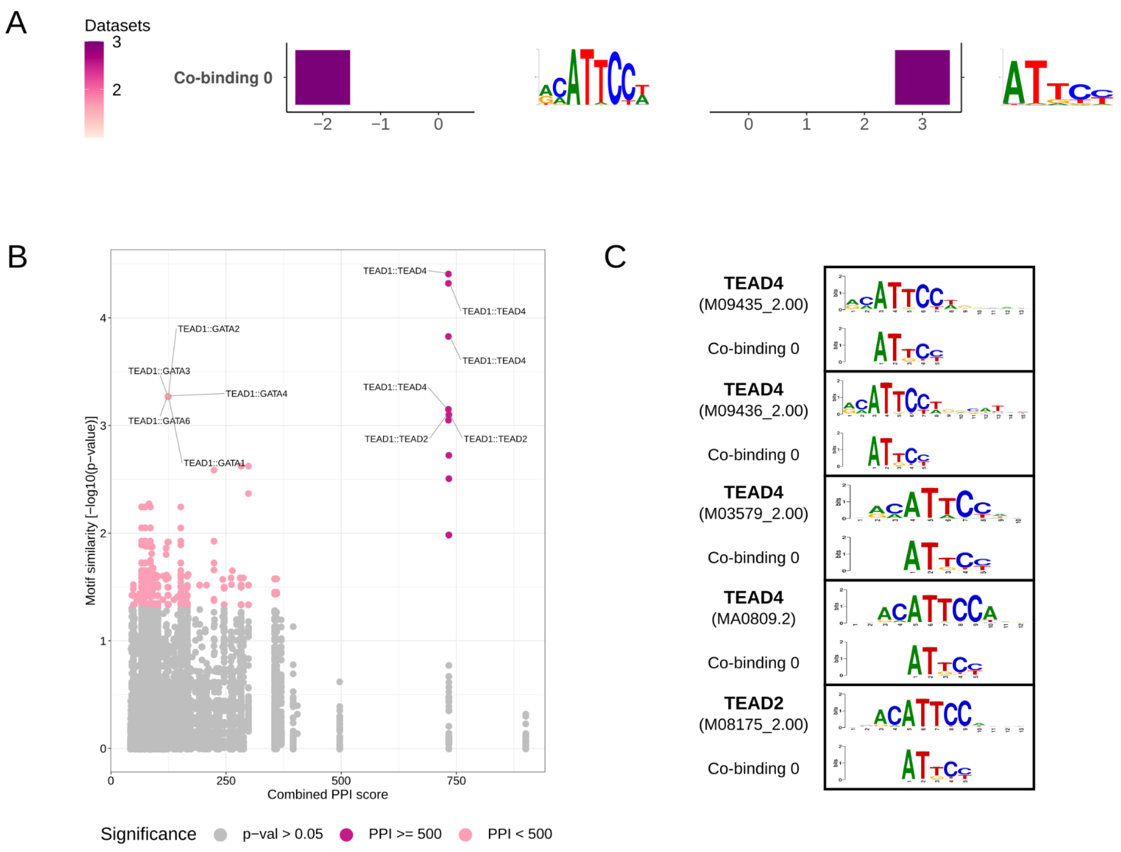


***Figure S9.*** **TEAD1 COBIND results.** COBIND predicts TEAD1 to homodimerize since the predicted motif is similar to the known binding motif associated with TEAD TFs, and TEAD1::TEAD1 obtains the highest combined PPI score. (**A**) The same co-binding motif is identified upstream (3 bp spacing) and downstream (2 bp spacing) of the anchor motif in the human and mouse genomes. (**B**) The co-binding motif has the highest similarity and PPI score (among mouse proteins) with TEAD proteins. (**C**) Discovered co-binding motifs match best with TEAD4 and TEAD2 TF motifs.

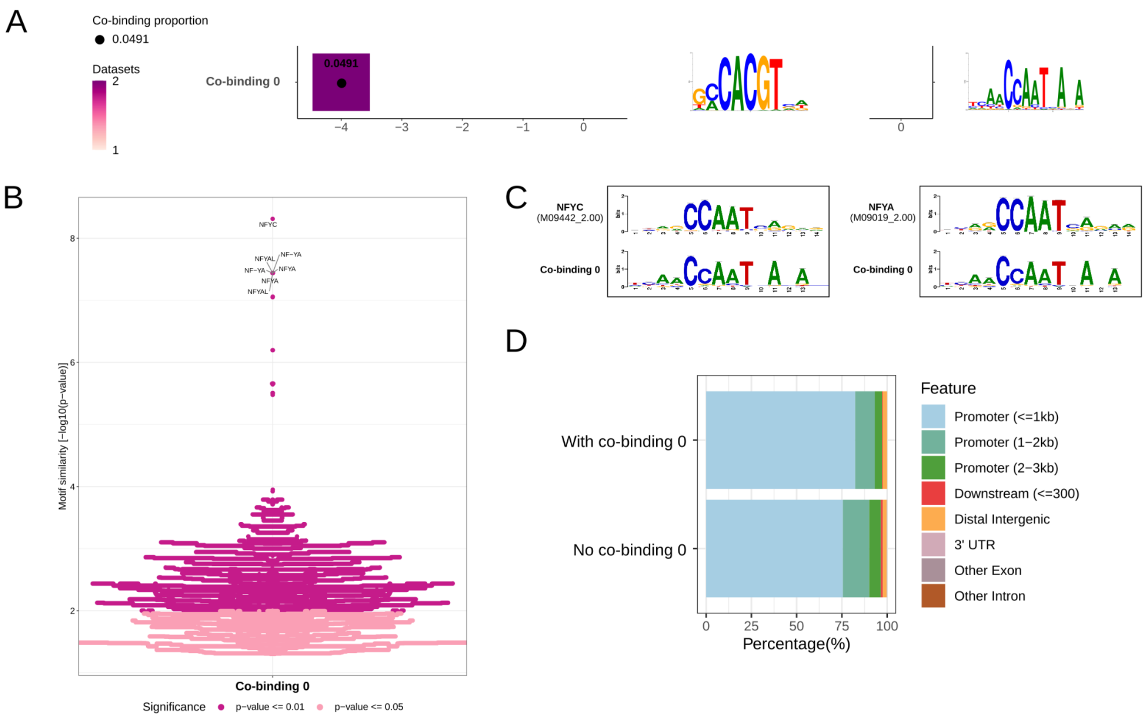


***Figure S10.*** **HY5 COBIND results.** (**A**) COBIND predicts a co-binding motif 4 bp upstream of the anchor HY5 TFBSs in the *Arabidopsis thaliana* genome. (**B**) The co-binder motif is similar to the motif bound by NF-Y family proteins. (**C**) The top matching motifs associate with the NF-YC and NF-YA binding profiles, but they are not known to interact physically with HY5 from the PPI data. (**D**) Most regions predicted to contain the co-binding pattern localize in promoter regions.

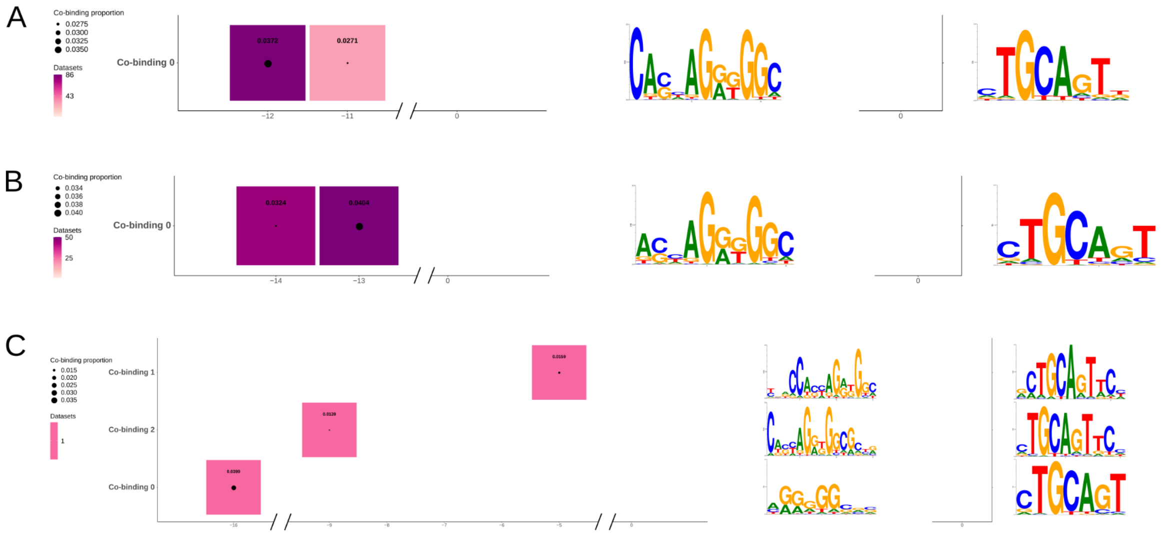


***Figure S11.*** **CTCF COBIND results across species.** (**A**) Mouse datasets of CTCF anchor TFBSs revealed two variants of the extended motif. (**B**) Human datasets of CTCF anchor TFBSs revealed the same motif but with different spacing patterns. (**C**) Zebrafish datasets of CTCF anchor TFBSs revealed the extended motif with variations within the corresponding anchor TFBSs.

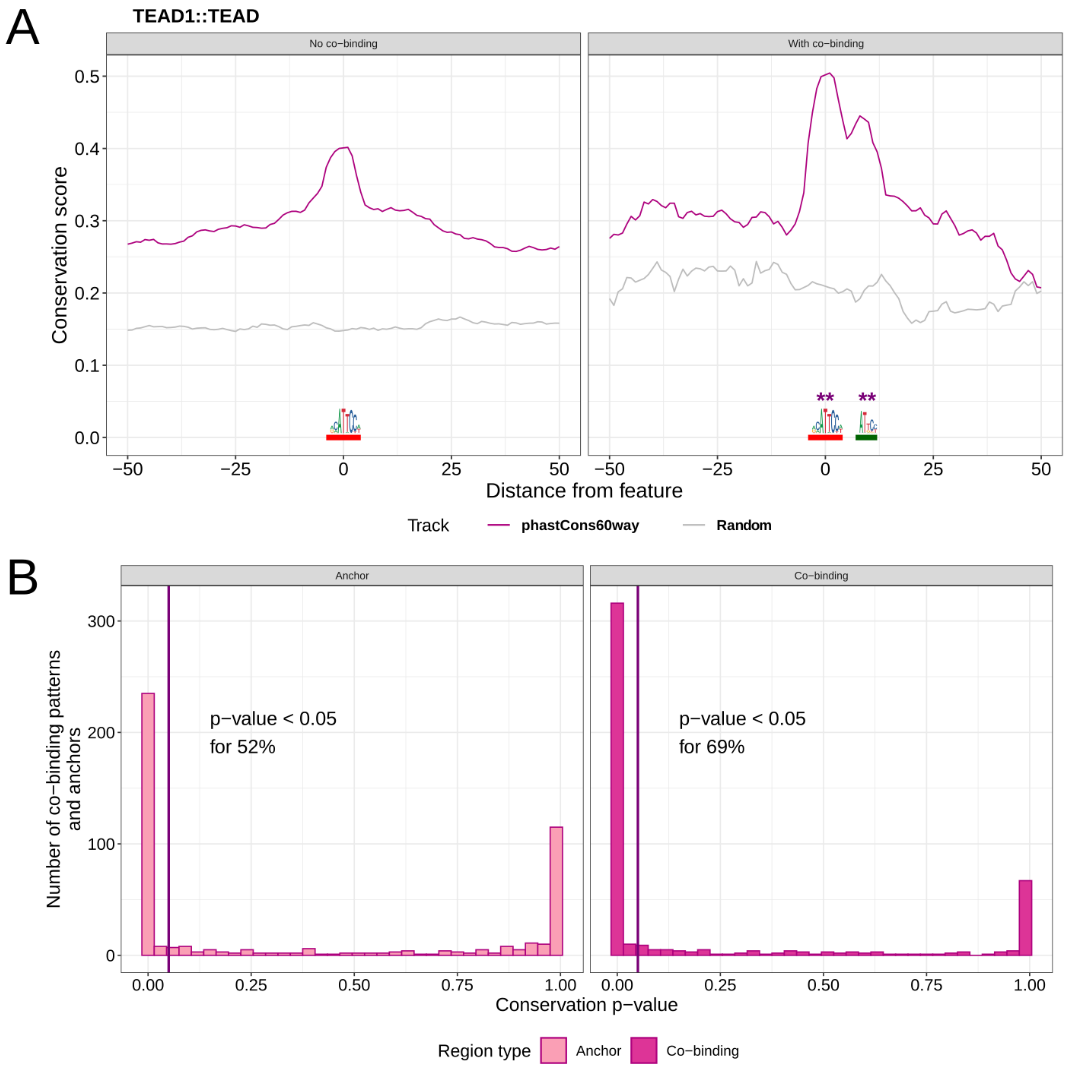


***Figure S12.*** **Analysis of the evolutionary conservation of co-binding patterns discovered by COBIND in the mouse genome.** (**A**) Comparison between vertebrate evolutionary conservation of genomic regions where COBIND predicted a co-binding pattern (right) or not (left). The purple lines provide the mean conservation score across the regions considered. The gray lines provide the mean conservation score across the same number of random genomic regions in the mouse genome. ** indicates a Wilcoxon test p-value < 0.001. The anchor TEAD1 motif is underlined in red, and the co-binding motif is underlined in green. (**B**) We compared the evolutionary conservation scores at the anchors' motifs in genomic regions harboring a co-binding pattern or not. The left panel represents the histogram of the number of co-binding patterns where the evolutionary conservation was higher (52%) in genomic regions harboring a co-binding pattern. The right panel represents the corresponding histogram when comparing genomic locations of the co-binding motif (69% of all co-binding patterns).

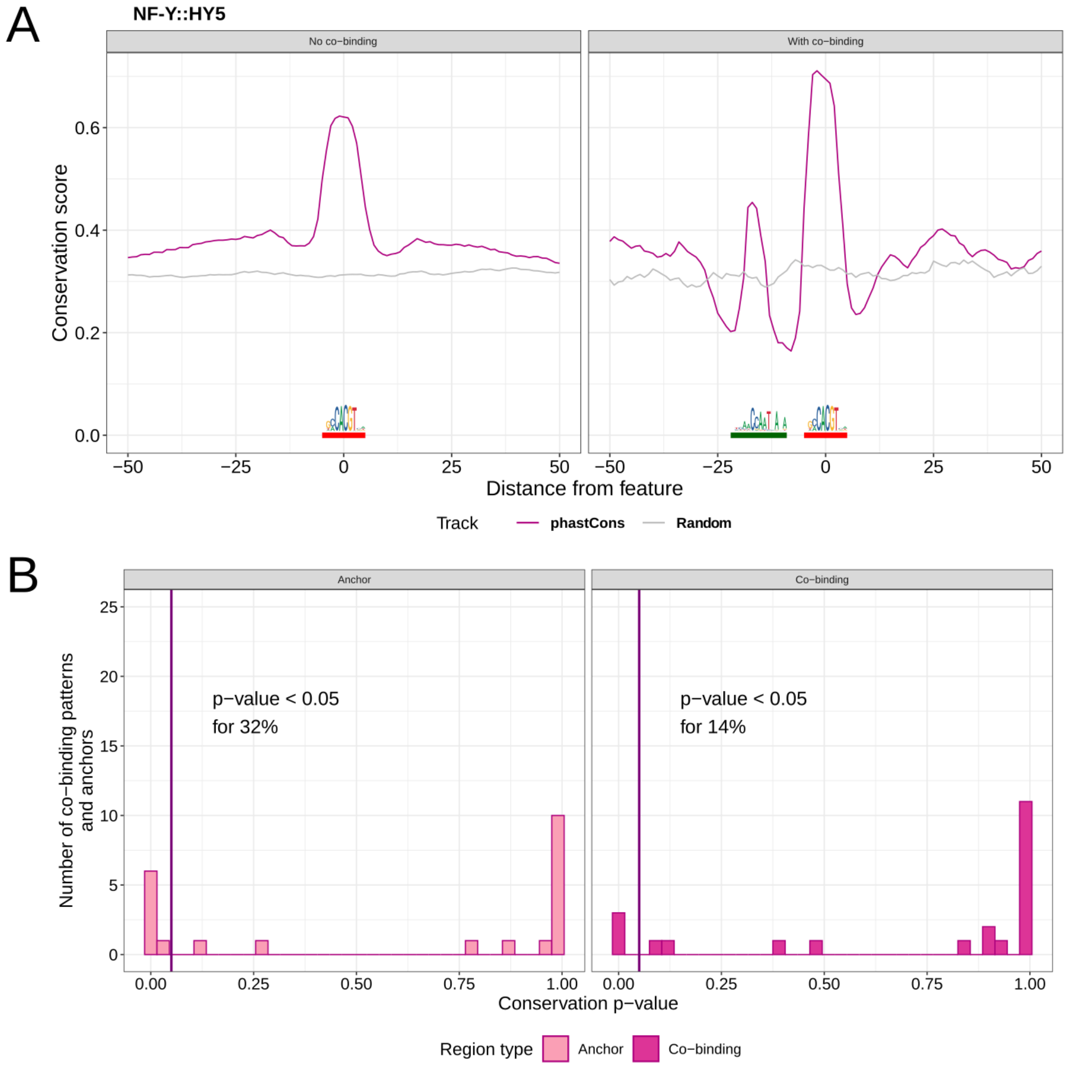


***Figure S13.*** **Analysis of the evolutionary conservation of co-binding patterns discovered by COBIND in the *A. thaliana* genome.** (**A**) Comparison between plant evolutionary conservation of genomic regions where COBIND predicted a co-binding pattern (right) or not (left). The purple lines provide the mean conservation score across the regions considered. The gray lines provide the mean conservation score across the same number of random genomic regions in the *A. thaliana* genome. The anchor HY5 is underlined in red, and the co-binding motif is underlined in green. (**B**) We compared the evolutionary conservation scores at the anchors' motifs in genomic regions harboring a co-binding pattern or not. The left panel represents the histogram of the number of co-binding patterns where evolutionary conservation was higher (32%) in genomic regions harboring a co-binding pattern. The right panel represents the corresponding histogram when comparing genomic locations of the co-binding motif (14% of all co-binding patterns).

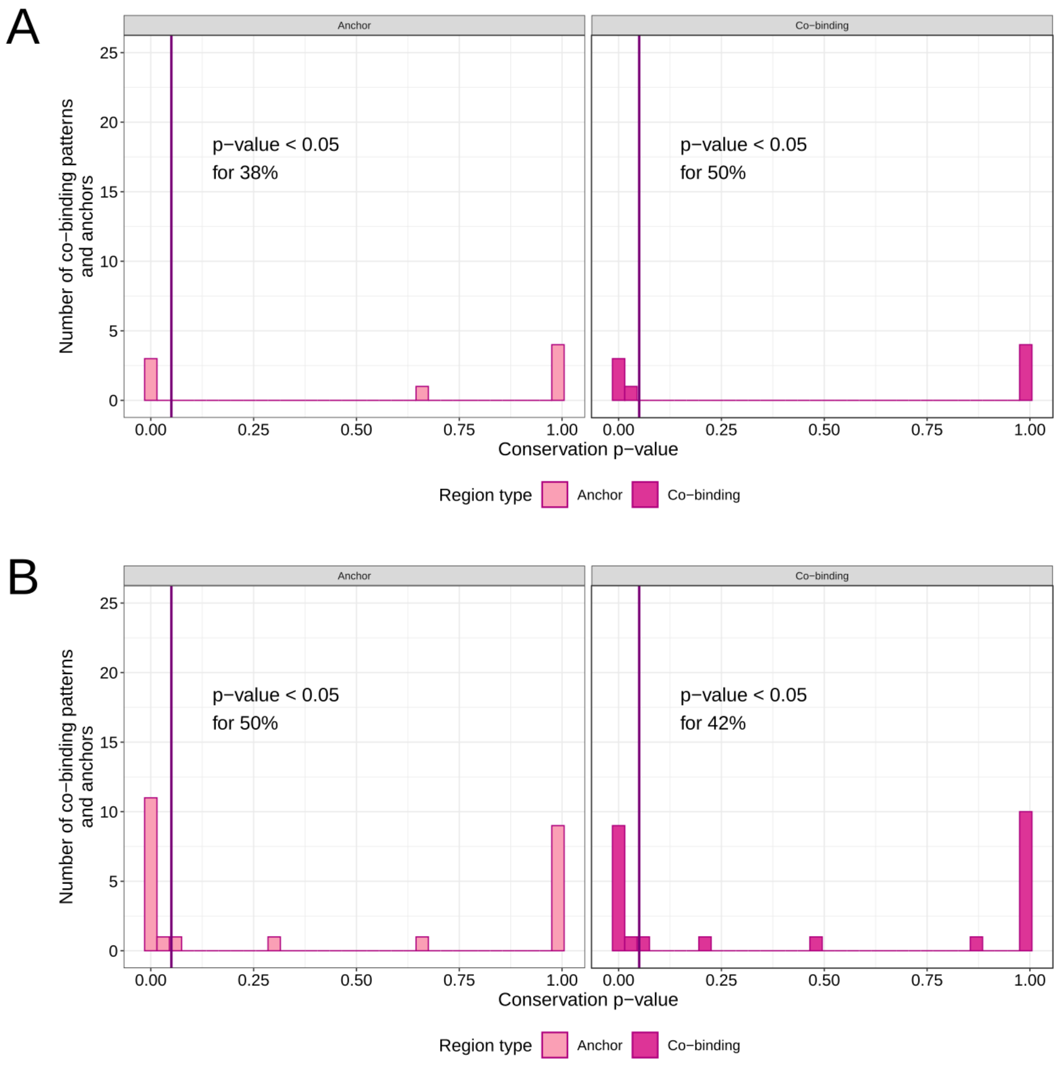


***Figure S14.*** **Analysis of the evolutionary conservation of co-binding patterns in the *C. elegans* (A) and *D. melanogaster* (B) genomes discovered by COBIND.** We compared the evolutionary conservation scores at the anchors' motifs in genomic regions harboring a co-binding pattern or not. The left panels in A and B represent the histograms of the number of co-binding patterns where evolutionary conservation was higher in genomic regions harboring a co-binding pattern (38% and 50%, respectively). The right panel represents the corresponding histogram when comparing genomic locations of the co-binding motif (50% and 42% of all co-binding patterns in *C. elegans* and *D. melanogaster*, respectively).

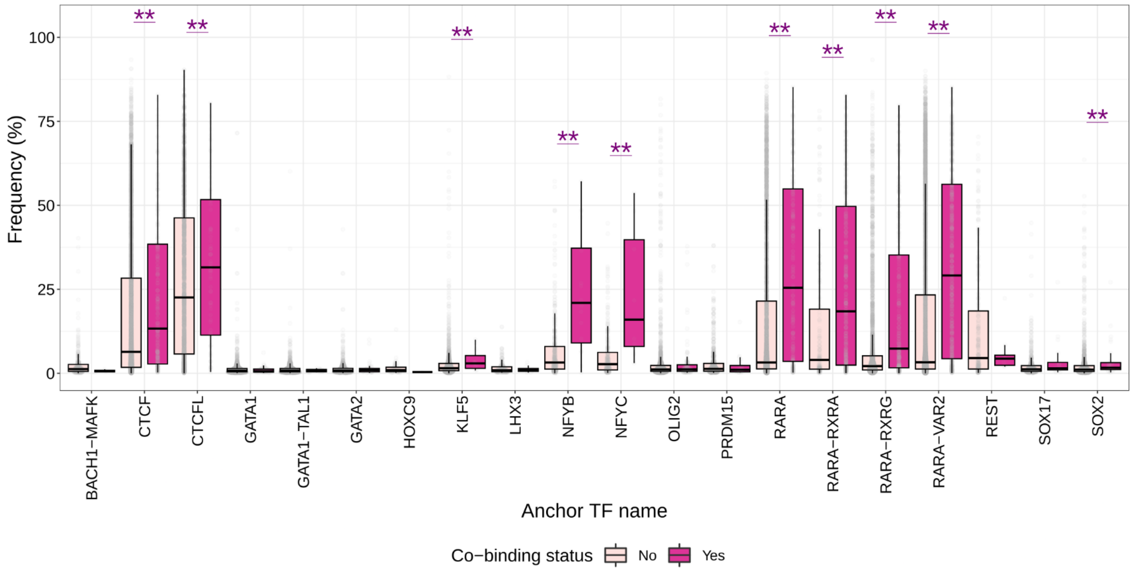


***Figure S15.*** **SMF data analysis across datasets.** Boxplots of the distribution of co-occupied molecules ("Anchor + Co-binding") overlapping regions with (dark pink) and without (light pink) a predicted co-binding pattern by COBIND when considering different anchor TFBSs. ** represents a Wilcoxon test p-value < 0.05.

### Supplementary Tables

***Table S1.*** **Parameters used when running the external tools within the COBIND pipeline.**

| Tool name/ Parameter name | Purpose | Value |
|---|---|---|
| Default COBIND parameters | | |
| Flank length *n* (base pairs) | Generating regions flanking the input anchors | `30` |
| Number of components *k* | A vector of *k* number of components to run NMF with | `[3, 6]` |
| Gini coefficient | A value from 0 to 1 used to filter discovered motifs | `0.5` |
| RSAT *matrix-clustering* | Clustering of discovered co-binder patterns | `-v 2 -calc sum -metric_build_tree cor -lth w 3 -lth cor 0.7 - lth Ncor 0.4` |
| | Clustering of associated anchor motifs | `-v 2 -calc sum -metric_build_tree cor -lth w 3 -lth cor 0.75 -lth Ncor 0.55` |
| Motif comparison | | |
| Tomtom v4.11.4 | Comparing *de novo* discovered motifs with inserted true motif (synthetic data analysis) or with known motif databases (TF inference through PPI analysis) | `-dist kullback -motif-pseudo 0.1 -min-overlap 1 -thresh 1 -eps -no-ssc` |

***Table S2.*** **UniBind 2021 robust collection data was used in this study**.

| Species name | Genome assembly | Number of files with TFBSs | Number of unique TFs | Number of unique TF motifs | Number of unique cell lines/tissues | Number of unique biological conditions |
|---|---|---|---|---|---|---|
| ***Arabidopsis thaliana*** | TAIR10 | 59 | 27 | 27 | 17 | 33 |
| ***Caenorhabditis elegans*** | WBcel235 | 59 | 19 | 19 | 11 | 41 |
| ***Danio rerio*** | GRCz11 | 6 | 4 | 5 | 3 | 3 |
| ***Drosophila melanogaster*** | dm6 | 73 | 16 | 16 | 24 | 36 |
| ***Homo sapiens*** | hg38 | 4,003 | 253 | 313 | 483 | 890 |
| ***Mus musculus*** | mm10 | 4,523 | 260 | 322 | 483 | 1,050 |
| ***Rattus norvegicus*** | Rnor 6.0 | 69 | 12 | 21 | 20 | 8 |

***Table S3.*** **Overview of tools for motif discovery and/or the investigation of TF cooperation.**

| | **COBIND** | **MEME** | **RSAT peak-motifs (*oligos, positions, dyads*)** | **SpaMo** | **TACO** | **iTFs** | **MCOT** |
|---|---|---|---|---|---|---|---|
| ***Year*** | ***2021*** | *2011* | *2011* | *2011* | *2014* | *2013* | *2019* |
| ***PubMed ID*** | ***This paper*** | *21486936* | *22156162*, *22836136* | *21602262* | *24640962* | *23847101* | *31750523* |
| **Input data** | Anchor regions in BED format (e. g., TFBSs) | Genomic sequences in fasta format | Genomic sequences in fasta format | Genomic sequences in fasta format, known primary (anchor) motif, motif library | ChIP-seq or DNase-seq peaks/regions, motif library | ChIP-seq peaks, motif library, TF pairs with known PPI | ChIP-seq peaks in fasta format, known primary (anchor) motif, motif library |
| **Anchor/Primary motif** | Anchor regions is input, but motif reflecting the anchor binding is not needed | MEME does not discover partner motifs. The assumption is that only one motif along the sequences is present and it is being discovered *de novo* in the input sequences | Similarly to MEME, peak-motifs tool is not intended for co-binding motif discovery. The algorithm discovers motif *de novo* using global over-representation (oligos) or positional bias (positions) or searching for two over-represented oligos (dyads) | Matrix of the anchor motif must be known and provided to the tool | Searches for matches in the provided motif library. Then anchor-co-binder pairs of different TFs are statistically tested | Genomic sites where a pair of TFs co-bind are predicted. Orientation and spacing bias is evaluated later with statistical tests | Motifs are searched *de novo* using HOMER. Motif is registered when there is a significant match with the "ChIP'ed" TF via Tomtom analysis |
| **Discovery of co-binder motifs** | Co-binders motifs upstream and downstream of the anchors are *de novo* discovered using non-negative matrix factorization (NMF). Each dataset is analyzed individually | | | Searches for matches in the provided motif library. Consensus sequences of a match and sequences for each match are returned | | | Searches for matches in the provided motif library. More specific matches can be achieved if ChIP-seq data in the same cell line/condition is available |

*(Continued in the next page)*

| | COBIND | MEME | RSAT peak-motifs (*oligos*, *positions*, *dyads*) | SpaMo | TACO | iTFs | MCOT |
|---|---|---|---|---|---|---|---|
| **Number of datasets** | Motifs are discovered in individual datasets, but co-binders can be summarized over groups of multiple datasets | Motifs are discovered in individual datasets | Motifs are discovered in individual datasets | Each dataset is analyzed individually | More than one dataset is required | Single dataset is enough | Single dataset is enough |
| **Summary of spacers and/or discovery of overlapping motifs** | Spacers are summarized and if overlap resulted in partial motif, such motif discovery is possible | - | Positional distribution is indicated, but post-processing would be needed to interpret it as a spacer. For dyads additional interpretation would be needed to distinguish anchor and co-binder | Spacer | Spacer and overlap | Spacer (categorized into ranges) | Spacer and overlap. A range of possible spacers should be defined by the user, default minimal 0 bp, maximum 29 bp |
| **Output** | Summary of spacings and locations of co-binders relative to the anchor motif together with sets of genomic regions with/without co-binders | Discovered motif logos and matrices together with discovery summary and combined discovered matrix | Discovered motif logos and matrices. Discovery summary with indicated motif positions in the set of sequences | Sequences where each motif from the library was found, results summary in html and tab-separated table formats | Dimer motifs, summary tables and genomic locations of cooperation events | Summary table | Anchor-co-binder pairs, spacing, regions |
| **Applications** | *De novo* discovery of TF co-binding motifs; discovery of extended TF motifs; summary of discovery, including spacings between reference and discovered motifs; other sequence pattern discoveries | *De novo* TF motif discovery | *De novo* TF motif discovery; differential motif discovery between two conditions | Identification of co-binding TFs and TF cooperativity analysis | Identification of co-binding TF pairs in DNase-seq and ChIP-seq datasets | Identification of co-binding TFs and TF cooperativity analysis | Identification of the co-binder motifs, analysis and summary of TF cooperativity |
| **Availability** | Command line | Web server, command line | Web server, command line | Web server, command line | Command line | Web server unavailable as of 2022 | Web server, command line |

*(Continued in the next page)*

| | COBIND | MEME | RSAT peak-motifs (*oligos*, *positions*, *dyads*) | SpaMo | TACO | iTFs | MCOT |
|---|---|---|---|---|---|---|---|
| **Scalability** | Parallelization over datasets and within the same dataset for motif discovery is possible through Snakemake | One command call for one dataset | One command call for one dataset | One command call for one dataset | Unknown | Unknown | One command call for one dataset |
| **Comments** | It is recommended to have at least 1000 regions in the input file. If less are available, Kim&Park and Gini thresholds can be adjusted | The tool does not perform any specific co-binding related analysis. It is recommended to input sequences of at least 100 bp in length | Oligos and Positions modes do not perform any specific co-binding related analysis. Dyad analysis could be seen as anchor-co-binder pair discovery, but is limited to simpler motif discovery | Highly dependent on the input motif library and no motif matrices for the co-binders are built and returned | Better predictions are achieved when there is cell type specific information. The tool was not included in the analysis due to code-running issues | Documentation, besides the paper, is not available. The tool was not included in the analysis due to code unavailability | The tool was not included in the analysis due to code-running issues |

**TF** - transcription factor

**TFBSs** - transcription factor binding sites
