## Additional File 2 for "Identification of transcription factor co-binding patterns with non-negative matrix factorization"

#### MATERIALS & METHODS

We downloaded intron coordinates for the human genome (hg38) using the UCSC Table Browser (1). We extracted the first two nucleotides of the exons flanking the introns. Depending on the direction of transcription, we defined these regions as *donors* or *acceptors*. We created two BED-formatted files - one for the donor and one for the acceptor sites, which we used as independent input for COBIND (default parameters).

#### RESULTS

We run COBIND on human donor and acceptor sites to see if COBIND discovers the expected splicing motifs (2). The acceptor motif consensus in different species is GT, and the consensus motif for donor sites is CAG (2, 3).

We anchored the COBIND analyses at the two exonic nucleotides flanking the introns. Thus, we analyzed the flanking regions upstream and downstream from the exon-intron borders. One flank is in the intron and another is in the exon (Materials and Methods above). COBIND revealed two patterns in the donor sites, where both have the consensus donor GT motif in around 7% of the input donor regions (Figure S1A). This consensus motif downstream of the anchor indicates the intronic flank. Moreover, the recovered anchor motif for the GTA motif (Co-binding pattern 0 in Figure S1A) contained high IC for AG, meaning that 4.44% of all input donor sites include AG as the last two exon nucleotides. When analyzing the acceptor sites, we found one DNA pattern corresponding to the consensus acceptor motif CAG in 1.35 % of input sequences (Figure S1B).

These analyses indicate that COBIND recovers the expected splicing signal motifs.

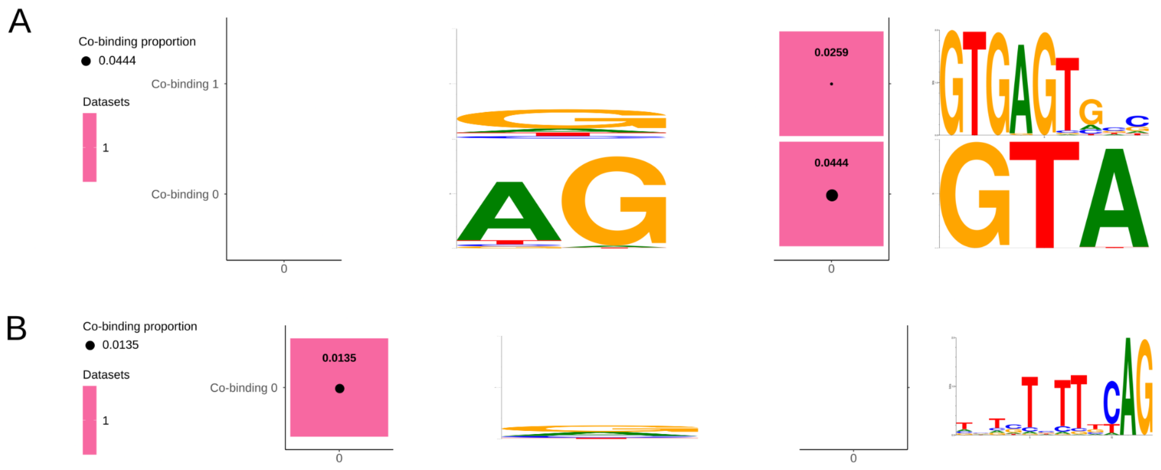


***Figure S1. COBIND*** **recovers splicing signal motifs at the intron-exon boundaries for donor (A) and acceptor (B) sites.**

### REFERENCES


1. Karolchik,D., Hinrichs,A.S., Furey,T.S., Roskin,K.M., Sugnet,C.W., Haussler,D. and Kent,W.J. (2004) The UCSC Table Browser data retrieval tool. *Nucleic Acids Res.*, **32**, D493–6.
2. Li,Y., Xu,Y. and Ma,Z. (2017) Comparative analysis of the Exon-intron structure in eukaryotic genomes. *Yangtze med.*, **01**, 50–64.
3. Mishra,A., Siwach,P., Misra,P., Dhiman,S., Pandey,A.K., Srivastava,P. and Jayaram,B. (2021) Intron exon boundary junctions in human genome have in-built unique structural and energetic signals. *Nucleic Acids Res.*, **49**, 2674–2683.
